# Depleting cationic lipids involved in antimicrobial resistance drives adaptive lipid remodeling in *Enterococcus faecalis*

**DOI:** 10.1101/2022.11.04.515160

**Authors:** Rafi Rashid, Zeus Jaren Nair, Dominic Ming Hao Chia, Kelvin Kian Long Chong, Amaury Cazenave Gassiot, Stewart A. Morley, Doug K. Allen, Swaine L. Chen, Shu Sin Chng, Markus R. Wenk, Kimberly A. Kline

**Author notes:** Contributed equally (Rafi Rashid, Zeus Jaren Nair). **Author Contributions**Conceptualization: RR, ZJN, KAK Formal analysis: RR, ZJN, DMHC, KKLC, ACG, SLC, SSC Funding acquisition: MRW, KAK Investigation: RR, ZJN, DMHC, KKLC, ACG, SLC Methodology: RR, ZJN, DMHC, KKLC, ACG, SLC, SSC, SAM, DKA Project administration: RR, ZJN, KAK Supervision: RR, MRW, KAK Writing – original draft: RR, ZJN, KAK Writing – review & editing: RR, ZJN, DMHC, KKLC, ACG, SLC, SSC, MRW, KAK.

## Abstract

The bacterial cell membrane is an interface for cell envelope synthesis, protein secretion, virulence factor assembly and a target for host cationic antimicrobial peptides (CAMPs). To resist CAMP killing, several Gram-positive pathogens encode the multiple peptide resistance factor (MprF) enzyme that covalently attaches cationic amino acids to anionic phospholipids in the cell membrane. While *E. faecalis* encodes two *mprF* paralogs, MprF2 plays a dominant role in conferring resistance to killing by the CAMP human β-defensin 2 (hBD-2) in *E. faecalis* strain OG1RF. The goal of the current study is to understand the broader lipidomic and functional roles of *E. faecalis mprF*. We analyzed the lipid profiles of parental wild type and *mprF* mutant strains and show that while ∆*mprF2* and ∆*mprF1* ∆*mprF2* mutants completely lacked cationic lysyl-phosphatidylglycerol (L-PG), the ∆*mprF1* mutant synthesized ∼70% of L-PG compared to the parent. Unexpectedly, we also observed a significant reduction of PG in ∆*mprF2* and ∆*mprF1* ∆*mprF2*. In the *mprF* mutants, particularly ∆*mprF1* ∆*mprF2*, the decrease in L-PG and PG is compensated by an increase in the phosphorus-containing lipid, GPDGDAG, and D-ala-GPDGDAG. These changes were accompanied by a downregulation of *de novo* fatty acid biosynthesis and an accumulation of long-chain acyl-acyl carrier proteins (long-chain acyl-ACPs), suggesting that the suppression of fatty acid biosynthesis was mediated by the transcriptional repressor FabT. Growth in chemically defined media lacking fatty acids revealed severe growth defects in the ∆*mprF1* ∆*mprF2* mutant strain, but not the single mutants, which was partially rescued through supplementation with palmitic and stearic acids. Changes in lipid homeostasis correlated with lower membrane fluidity, impaired protein secretion, and increased biofilm formation in both ∆*mprF2* and ∆*mprF1* ∆*mprF2*, compared to wild type and ∆*mprF1*. Collectively, our findings reveal a previously unappreciated role for *mprF* in global lipid regulation and cellular physiology, which could facilitate the development of novel therapeutics targeting MprF.

**Significance Statement:** The cell membrane plays a pivotal role in protecting bacteria against external threats, such as antibiotics. Cationic phospholipids such as lysyl-phosphatidyglycerol (L-PG) resist the action of cationic antimicrobial peptides through electrostatic repulsion. Here we demonstrate that L-PG depletion has several unexpected consequences in *Enterococcus faecalis*, including a reduction of phosphatidylglycerol (PG), enrichment of a phosphorus-containing lipid, reduced fatty acid synthesis accompanied by an accumulation of long-chain acyl-acyl carrier proteins (long chain acyl-ACPs), lower membrane fluidity, and impaired secretion. These changes are not deleterious to the organism as long as exogenous fatty acids are available for uptake from the culture medium. Our findings suggest an adaptive mechanism involving compensatory changes across the entire lipidome upon removal of a single phospholipid modification. Such adaptations must be considered when devising antimicrobial strategies that target membrane lipids.

## Introduction

*Enterococcus faecalis* is a Gram-positive commensal bacterium that naturally inhabits the harsh environment of the human gastrointestinal tract. The Enterococci are amongst the most clinically significant nosocomial pathogens and cause a variety of opportunistic infections in susceptible individuals, including endocarditis, urinary tract infections, bacteremia, and wound infections (1). Complicating the management of these infections is the fact that Enterococci are naturally resistant to conventional antibiotics like aminoglycosides; have rapidly evolved resistance to other drugs like chloramphenicol, erythromycin, tetracyclines, and vancomycin (2); and readily form biofilm during infection, conferring phenotypic antibiotic tolerance on these organisms (3).

For Enterococci to successfully colonize their human hosts, they must resist the bactericidal effect of cationic antimicrobial peptides (CAMPs), amphipathic molecules that are part of the innate immune response against microbes during infection (4). CAMPs are produced by a wide range of organisms, including bacteria, fungi, plants, and animals, and exert their antibacterial effects via disruption of the cell membrane (5). In humans, CAMPs are divided into two main classes, the defensins, e.g., human neutrophil peptides (HNP) and human β-defensins (hBD), and the cathelicidins, e.g., LL-37 (6, 7). The positively charged moiety of CAMPs binds to negatively charged membrane phospholipid headgroups, whereas the hydrophobic moiety binds to hydrophobic regions created by phospholipid acyl chains. Bacterial cell death results from physical disruption of the membrane, for which three different models (barrel-stave, toroidal, and carpet) have been described (reviewed in (8)). More recently, a new mechanism of disruption in which CAMPs delocalize membrane lipids and proteins has been reported for some bacteria (8, 9).

In *E. faecalis*, CAMPs are thought to target phosphatidylglycerol (PG), an anionic phospholipid which constitutes a major portion of the cell membrane (10, 11). To evade CAMP killing, Gram-positive bacteria have evolved mechanisms to modify their anionic phospholipids (9, 12-14). For example, esterification of PG with the cationic amino acid lysine enables *Staphylococcus aureus* to resist the killing action of CAMPs through electrostatic repulsion. This lysinylation of PG changes its net charge from negative to positive. Because this modification confers resistance to multiple CAMPs, the gene responsible was named *mprF* for multiple peptide resistance factor. MprF contains two domains that perform distinct enzymatic functions: a synthase domain responsible for modifying PG with amino acids (e.g., lysine), and a flippase domain responsible for transferring modified PG from the inner leaflet to the outer leaflet of the cytoplasmic membrane (15). MprF has since been shown to confer protection against CAMPs in several other Gram-positive organisms including *Bacillus anthracis, Bacillus subtilis, E. faecalis, Listeria monocytogenes, Mycobacterium tuberculosis*, as well as in Gram-negative *Pseudomonas aeruginosa* (9, 12-14, 16-18).

We previously showed that MprF2 confers resistance to killing by the CAMP human β-defensin 2 (hBD-2) in *E. faecalis* strain OG1RF (9). Our findings are consistent with an earlier study which showed that deleting the *mprF2* gene in *E. faecalis* 12030 resulted in loss of L-PG and higher sensitivity to the CAMPs colistin, nisin, HBD-3, and polymyxin B (14). In OG1RF where selected phospholipid synthesis genes (including *mprF2*) were deleted, exogenous fatty acid supplementation resulted in tolerance to the cationic antibiotic daptomycin and changes in lipid composition (19). Mutations in genes encoding the LiaFSR 3-component system and phospholipid metabolism enzymes accompanied alterations in phospholipid composition in daptomycin-resistant strains of *E. faecalis* (20, 21). Such adaptive lipid remodeling also occurs in other bacterial species in response to environmental changes (22) and phospholipase activity (23).

Our previous work and the above literature linking *mprF* genetic perturbation with cationic antimicrobial resistance and lipidomic alterations suggest a gap in knowledge about how *mprF* brings about such adaptive lipid remodeling. Thus, we sought to more thoroughly investigate the mechanism by which *mprF* affects lipid homeostasis in *E. faecalis*. Using lipidomic methods that we had previously established (24), we quantified lipids across various classes in wild type OG1RF and the ∆*mprF1*, ∆*mprF2*, and ∆*mprF1* ∆*mprF2* mutant strains. Coupled with transcriptomic analyses, gas chromatography analysis of total fatty acid methyl esters (GC-FAME), assays for membrane fluidity, secretion, biofilm formation, and CAMP susceptibility, we have discovered a previously unappreciated role for *mprF* in global lipid regulation and cellular physiology.

## Results

### *mprF* deletion alters levels of membrane phospholipids

We previously observed that human β-defensin 2 (hBD-2) binds *E. faecalis* more strongly at foci situated at or near the division septum in the ∆*mprF2* mutant than in the wild type, and that only MprF2, and not MprF1, protects against hBD-2-mediated killing (9). The ∆*mprF2* strain is also more susceptible to LL-37, a cathelicidin that is more potent than hBD2, compared to the ∆*mprF1* strain, and ∆*mprF1* ∆*mprF2* is significantly more susceptible to hBD-2 than ∆*mprF1* (but not ∆*mprF2*) at 50 µg/ml **(Fig. S1A, S1B)**. We hypothesized that differences in CAMP susceptibility between *mprF1* and *mprF2* mutants may be due to differences in levels of aminoacylated lipids. Previously, we performed untargeted lipidomics to identify the phospholipid and glycolipid repertoire of *E. faecalis* OG1RF, and multiple reaction monitoring (MRM) to quantify phosphatidylglycerol (PG) and lysyl-PG (L-PG) molecules in this strain and in two daptomycin-resistant strains (10). In the current study, we used the same methodology to compare the levels of L-PG in ∆*mprF1*, ∆*mprF2*, and ∆*mprF1* ∆*mprF2* mutants with the parental wild type strain. We found that ∆*mprF2* and ∆*mprF1* ∆*mprF2* cells had no detectable L-PG **(Fig. 1A, S1C)**, as previously described by Bao and colleagues (14), whereas total L-PG was present in the ∆*mprF1* mutant at 70% of the wild type level. Unlike previous thin layer chromatography (TLC) studies which lacked the sensitivity to discriminate between individual lipid molecules and quantify their levels, using LC-MS/MS we were able to observe that MprF1 also contributes to L-PG synthesis in *E. faecalis* OG1RF under these laboratory conditions.

**Figure 1.**
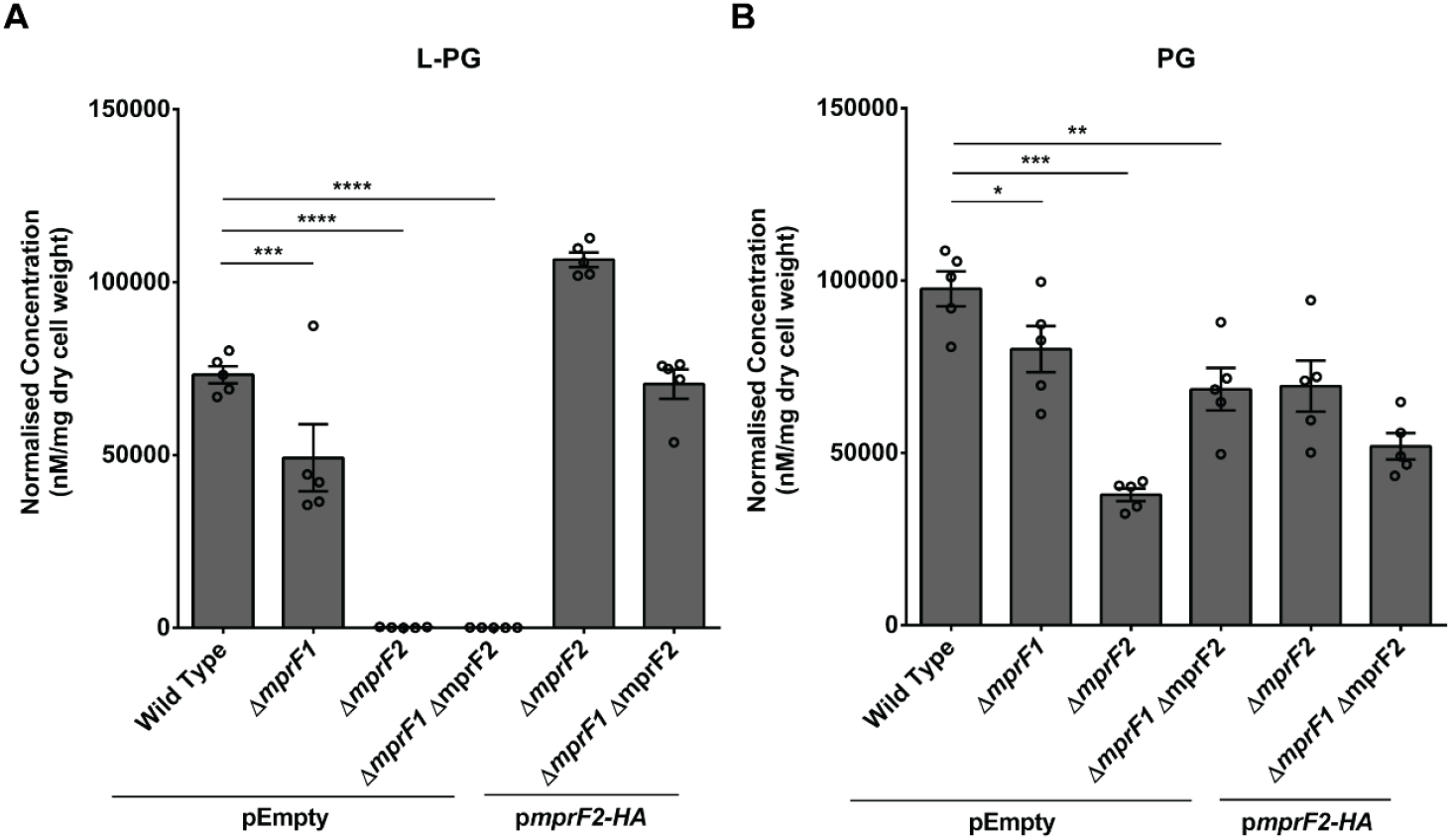
*mprF* contributes to L-PG and PG levels. Normalized amounts of total lysyl-phosphatidylglycerol (L-PG) **(A)** and total phosphatidylglycerol (PG) quantities **(B)** in *E. faecalis* wild type and *mprF* mutants are shown. These amounts were obtained by normalizing against internal standards and dry cell weight of the respective samples. *mprF2*-HA refers to *mprF2* with a hemagglutinin (HA) affinity tag on its C-terminal for ease of assessing expression. Data showing complementation with untagged *mprF2* can be found in **Fig. S1E-H**. Each bar represents the mean ± standard error of measurement calculated from 5 biological replicates, represented by each open circle. *, p<0.05; **, p<0.01; ***, p<0.001; ****, p<0.0001; Fisher’s LSD test for ANOVA.

To determine whether the above differences in L-PG levels were due to differential *mprF* gene expression, we performed quantitative reverse transcription polymerase chain reaction (RT-qPCR) of the *mprF* paralogs in the wild type. RT-qPCR revealed that *mprF2* expression is 200-fold higher than *mprF1* expression **(Fig. S2A)**. This differential expression would explain MprF1’s relatively minor contribution to L-PG production. We also observed that deleting either *mprF* paralog does not affect the expression of the remaining paralog **(Fig. S2B)**.

As L-PG synthesis requires PG as a substrate, we predicted that PG levels would be higher in mutants unable to convert a fraction of PG to L-PG. Unexpectedly, ∆*mprF2 and* ∆*mprF1* ∆*mprF2* mutants had significantly less PG than wild type. Moreover, similar to our observation for L-PG, ∆*mprF1* also had an intermediate amount of PG compared to wild type and the other two *mprF* mutants (**Fig. 1B, S1D**). The reduction in PG was unexpected because its biosynthetic gene (*pgsA*) is thought to be essential in *E. faecalis*, based on the fact that this gene is essential in many other bacterial species (25). However, our mutants do not show any growth defect **(Fig. S2C)**, change in viability **(Fig. S2D)**, or increased membrane permeability to propidium iodide **(Fig. S2E)**.

Complementing the ∆*mprF2* mutant with *mprF2* on a plasmid more than restored L-PG to wild type levels **(Fig. 1A, S3A)** and partially restored PG levels **(Fig. 1B, S3B)**, the latter possibly being the result of the complemented strain producing more L-PG than the wild type and therefore depleting the PG precursor. When the double mutant was complemented with the *mprF2* gene, L-PG was restored to wild type levels, whereas the level of PG was lower than in the uncomplemented double mutant **(Fig. 1A, 1B)**. To confirm that it is MprF2’s enzymatic activity in L-PG production (rather than another indirect effect of MprF2) that determines PG levels, we complemented the ∆*mprF2* mutant with inactive *mprF2* on a plasmid. This complementation failed to restore L-PG to wild type levels **(Fig. S1E, S1G)**. The catalytically inactive *mprF2* possesses mutations in residues coordinating its interaction with the aminoacyl moiety of lys-tRNA at similar locations (D739A, R742S) as previously described in *B. licheniformis* (26). Inactive *mprF2* complementation also failed to restore PG to levels achieved through complementation with native MprF2 **(Fig. S1F, S1H)**.

We also considered the individual effects of each paralog’s N-terminal flippase and C-terminal catalytic domains on L-PG and PG levels. Complementation with both domains of *mprF2* is required for L-PG production **(Fig. S3A)**. Restoration of L-PG and PG levels was not possible through complementation with either full length *mprF1* or individual domains of either paralog. **(Fig. S3A-D)**.

Collectively, our results suggest that deletion of *mprF1, mprF2*, or both genes causes more widespread changes in phospholipid homeostasis than previously appreciated.

### *mprF* deletion affects lipids in other classes

In light of reduced L-PG and PG in the *mprF* mutants and the fact that PG is an essential lipid in many Gram-positive bacteria (27-29), we hypothesized that the mutants might adjust the abundances of other lipid classes via adaptive lipidome remodeling (22) to compensate for the depletion of otherwise essential lipids. We first conducted one-dimension thin layer chromatography (1D-TLC) of lipid extracts of each strain, using 14C radiolabeling and iodine staining **(Fig. 2A)**. Six major spots were observed in the 1D-TLC which were identified as diglucosyl-diacylglycerol (DGDAG), L-PG, glycerophospho-diglucosyl-diacylglycerol (GPDGDAG), D-ala-GPDGDAG, PG and cardiolipin (CL) co-migrating in a single spot, and a spot of unknown identity containing phosphorus **(Fig. 2A)**. Identities of each spot were determined by a combination of 1- and 2-dimension TLC analyses with lipid standards, ninhydrin staining, mass spectrometry, and 32P and 14C radiolabeling as described in **Supplementary Text 1A**.

**Figure 2.**
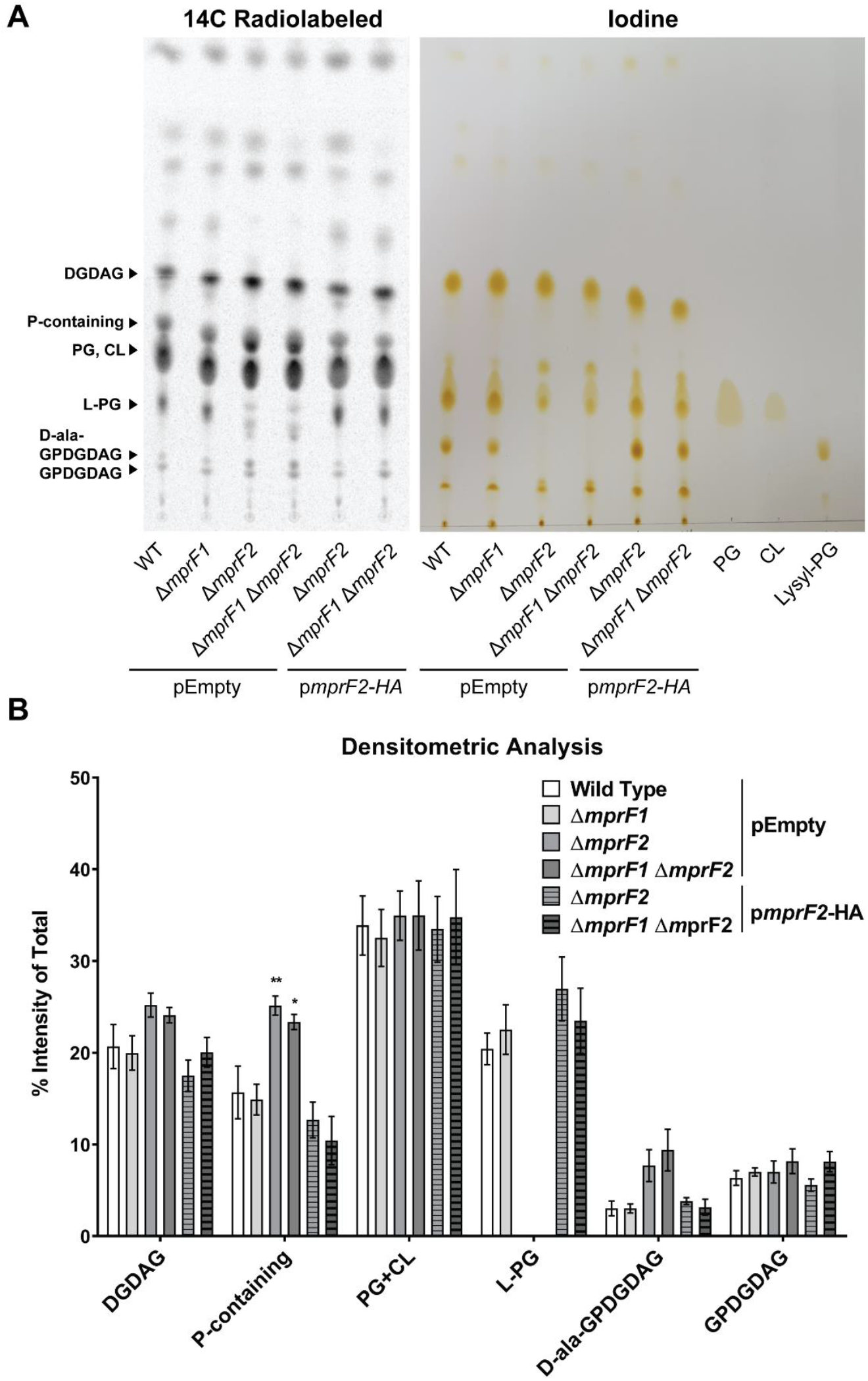
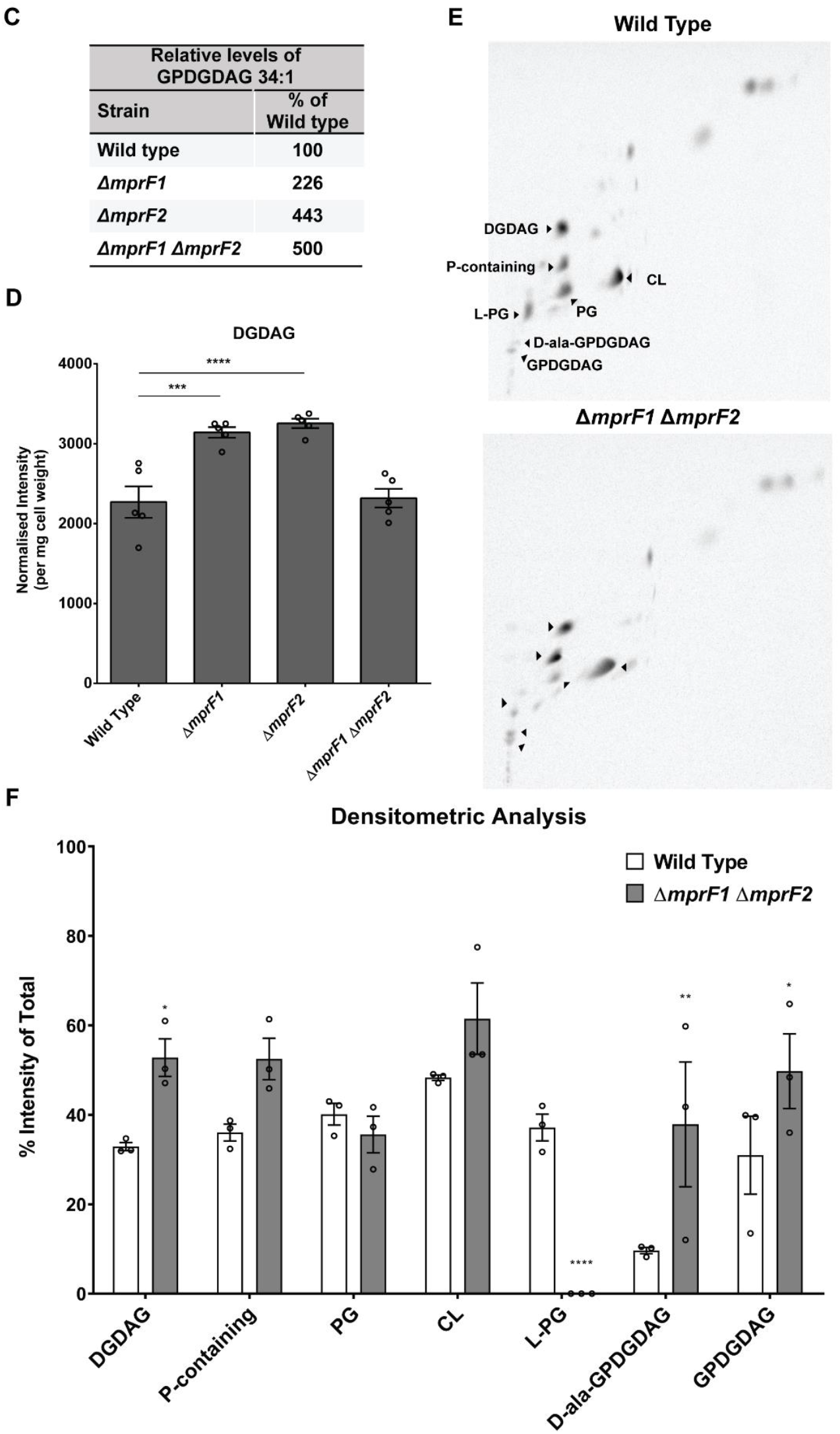
*mprF* deletion results in changes in other lipid classes. **(A)** Representative _14_C radiolabeled and iodine stained 1D-TLC of lipid extracts from the *mprF* mutants using chloroform: methanol: water (65:25:4). Standards were applied to the iodine-stained TLC plate to confirm the positions of PG, CL, and L-PG spots. Further identification of the other lipid spots can be found in the supplement (Supplementary Section 1 - TLC spot identification). **(B)** Densitometric analysis of _14_C radiolabeled 1D TLC spots. Each bar represents the mean ± standard error of measurement calculated from 4 biological replicates. **(C)** Semiquantitative analysis of dominant GPDGDAG species, GPDGDAG 34:1. **(D)** Semiquantitative quantification of DGDAG amounts across the *mprF* mutants. These amounts were obtained by normalizing against an external surrogate standard (MGDAG 34:1) and dry cell weights of the respective samples. Each bar represents the mean ± standard error of measurement calculated from 5 biological replicates. **(E)** Representative 2D-TLC of lipid extracts from wild type and ∆*mprF1* ∆*mprF2* for further spot separation to visualize changes in PG and CL. **(F)** Densitometric analysis of 14C radiolabeled 2D TLC spots. Each bar represents the mean ± standard error of measurement calculated from 3 biological replicates. Statistical comparisons made for mutants against wild type. *, p<0.05; **, p<0.01; ***, p<0.001; ****, p<0.0001; Fisher’s LSD test for ANOVA. Identities of these spots were determined through a combination of TLCs with lipid standards as well as mass spectrometry of the spots. Detailed information on how spot identities were assigned can be found in the supplement **(Supplementary Text 1A, Fig. S4**, and **Excel Table S1C)**.

Both the 14C radiolabeled and iodine-stained TLC plates indicate that L-PG is lower only in ∆*mprF2* and ∆*mprF1* ∆*mprF2* and that reduction was restored by complementation with p*mprF2*-HA **(Fig. 2A)**, findings which corroborate the quantification data obtained via LC-MS/MS-based multiple reaction monitoring **(Fig. 1A)**. We observed more intense spots of GPDGDAG and D-ala-GPDGDAG in these mutants compared to wild type, although the difference was not statistically significant. Combined PG and CL spot intensities were unchanged across the different mutants.

We also observed stronger spot intensities for the phosphorus-containing lipid of unknown identity in ∆*mprF2* and ∆*mprF1* ∆*mprF2* **(Fig. 2B)**. Although we have yet to identify this lipid, TLC analysis confirmed that the spot is a lipid (positive iodine staining and presence of fatty acyl product ions from MS2 analysis) that contain phosphorus (positive 32-P radiolabeling) but no amino groups (negative ninhydrin staining) **(Fig. 2B, S4C-E)**. Based on MS analysis, this unknown lipid spot contains 18 species that do not belong to any of the lipid classes we have previously studied (i.e. PG, LPG, Ala-PG, Arg-PG, DGDAG, MGDAG, GPDGDAG). MS-MS analysis of the precursor ions reveals that they contain fatty acyl chains and three common products ions (m/z: 379.1, 397.1, 415.1) that were consistent across all species, suggesting that these species belong to the same class **(Supplementary Excel Table S1C)**. Complementation with p*mprF2*-HA restored this lipid to wild type levels **(Fig. 2A)**.

As PG is a substrate for GPDGDAG synthesis (30), we asked whether GPDGDAG levels were significantly altered in mutants where PG was reduced. 14C-radiolabeled TLC revealed faint spots for GPDGDAG. Since it was not possible to discern any differences in GPDGDAG levels via TLC **(Fig. 2B)**, we performed a semi-quantitative mass spectrometric analysis on one particular GPDGDAG species, GPDGDAG 34:1 **(Fig. 2C)** that we previously determined to be the most abundant species within the class (10). By this latter analysis, ∆*mprF1* ∆*mprF2* showed a 5-fold increase in GPDGDAG 34:1 levels relative to wild type, with ∆*mprF1* and ∆*mprF2* showing 2- and 4-fold higher GPDGDAG 34:1 levels, respectively **(Fig. 2C)**. Absolute quantification via MRMs was not possible due the lack of a commercially available standard.

Since GPDGDAG 34:1 was higher in the *mprF* mutants, we considered whether the abundance of its downstream product, lipoteichoic acid (LTA) (which is poly-glycerophosphate polymerized onto a GPDGDAG membrane anchor) might be affected as well (31). However, immunoblots of whole cell lysates and supernatants from the wild type and *mprF* mutants revealed no difference in LTA levels in the whole cell lysates, and no shed LTA in the supernatants of any strain, suggesting that LTA levels remain unchanged in these mutant backgrounds **(Fig. S1I)**.

Due to the lack of a commercially available DGDAG standard, we performed semi-quantitative MRMs for DGDAG using a monoglucosyl-diacylglycerol (MGDAG) surrogate standard **(Fig. 2D)**. ∆*mprF1* and ∆*mprF2* had more total DGDAG than the wild type, while DGDAG in ∆*mprF1* ∆*mprF2* was at the wild type level **(Fig. 2D)**. The same trend was observed for most of the individual DGDAG species **(Fig. S1J)**.

To discriminate between individual PG and CL spot intensities, we performed 2D TLCs to further separate the co-migrating PG and CL spots in the first dimension **(Fig. 2E)**. We observed that the PG spot intensity was slightly lower in ∆*mprF1* ∆*mprF2* while CL appears unchanged **(Fig. 2F)**. Similar to the 1D-TLCs, L-PG appeared to be lower while the unknown phosphorus-containing lipid spot, together with the D-ala-GPDGDAG and GPDGDAG spots, appeared stronger in intensity in ∆*mprF1* ∆*mprF2* **(Fig. 2F)**. However, DGDAG was stained more intensely **(Fig. 2E, 2F)** unlike what was observed in the MS analysis **(Fig. 2C)**. This difference could be due to another 14C-radiolabeled compound co-migrating with DGDAG, resulting in more intensely stained 2D- and 1D-TLCs (though the staining intensity difference is not statistically significant) **(Fig. 2B, 2F)**. All of these observations suggest that in the *mprF* mutants, particularly ∆*mprF1* ∆*mprF2*, the decrease in L-PG and PG is compensated by an increase in the phosphorus-containing lipid, GPDGDAG, and D-ala-GPDGDAG.

### *mprF* deletion results in decreased expression of fatty acid synthesis genes

Since *mprF* deletion resulted in alterations in phospholipid and glycolipid levels, we hypothesized that these altered levels might be due to altered expression of genes involved in lipid metabolism. To test this hypothesis, we performed RNA sequencing to compare the gene expression profiles of the wild type and the *mprF* mutants. In total, 301 genes (14% of the genome) were differentially expressed between the wild type and ∆*mprF1* ∆*mprF2* **(Fig. S5A)**. Principal component analysis revealed that the transcriptomic profile of ∆*mprF2* clusters more closely to that of ∆*mprF1* ∆*mprF2*, suggesting that *mprF2* deletion is associated with more transcriptional changes than *mprF1* deletion **(Fig. S5B)**. A Venn diagram of genes differentially expressed in the *mprF* mutants relative to wild type indicates that *mprF1* and *mprF2* have individual effects that are amplified in the double mutant, from which we infer that the mutant phenotypes are not only due to epistasis (**Fig. S5C)**. We performed a statistical analysis of gene sets in all four strains and, consistent with our hypothesis, we found downregulation of nine genes associated with fatty acid metabolism and biosynthesis in ∆*mprF1* ∆*mprF2* **(Fig. 3A, B)**. Genes involved in the initiation, elongation, and termination phases of fatty acid synthesis *(accA*, accB, *accC, accD, fabD, fabF2, fabG3, fabK*, and *fabZ2)* were downregulated in the double mutant **(Fig. 3B, S5A)**. Using RT-qPCR, we confirmed that genes encoding components of the *de novo* fatty acid biosynthesis pathway were downregulated in the double mutant. By contrast, expression of the essential gene *pgsA*, which encodes an enzyme involved in PG synthesis, was slightly upregulated in the double mutant **(Fig. S5D)**, perhaps in an attempt to compensate for the decrease in the levels of PG.

**Figure 3.**
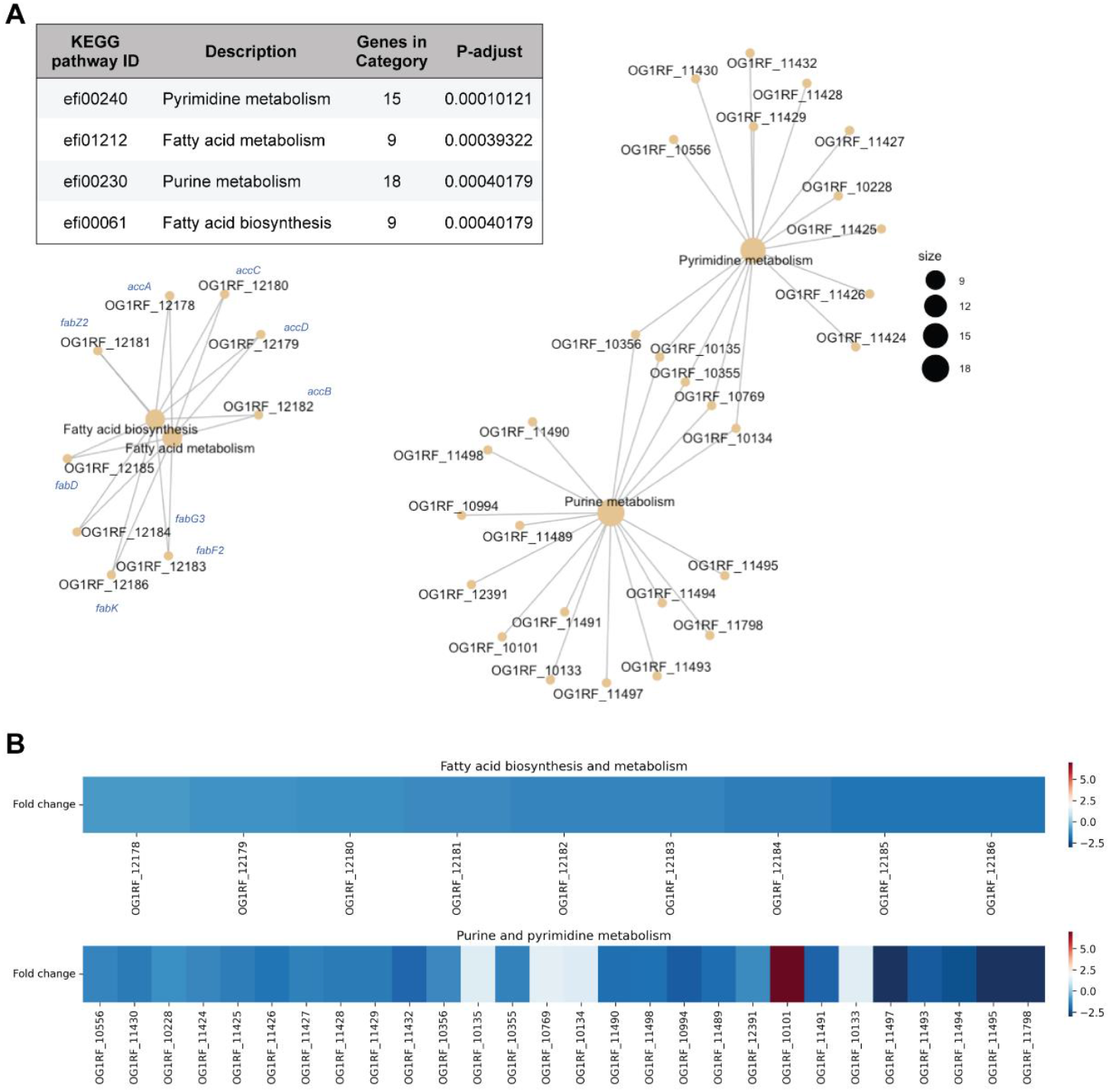
Loss of *mprF* leads to changes in regulation of fatty acid biosynthesis and nucleotide metabolism genes. **(A)** Gene set enrichment analysis comparing KEGG categories of differentially expressed genes between the parental wild type and *ΔmprF1 ΔmprF2*. The table shows categories that are differentially regulated and the plots showing interconnectivity between genes in each differentially regulated pathway. Node size corresponds to the number of enriched genes. **(B)** Heatmap showing differential gene expression based on the gene set enrichment analysis in **(A)**.

### Saturated fatty acids restore *mprF* growth in nutrient-limited media

Since *de novo* fatty acid biosynthesis is downregulated in ∆*mprF1* ∆*mprF2*, we speculated that this mutant might require exogenous fatty acids for growth and survival. *E. faecalis* can incorporate exogenous fatty acids into its membrane when grown in bovine heart infusion (BHI) culture medium (32). We performed a gas chromatography analysis of total fatty acid methyl esters (GC-FAME) on BHI, in which we detected only two fatty acids - palmitic (C_16:0_) and stearic acid (C_18:0_) **(Supplementary Excel Table S1D)**. We thus hypothesized that ∆*mprF1*∆*mprF2* takes up these fatty acids from BHI to counteract the downregulation of *de novo* fatty acid biosynthetic genes. As expected, we observed that growth of the double mutant was severely impaired in nutrient-limited, chemically defined media (CDM) lacking fatty acids, while growth of the single mutants was less severely impaired **(Fig. 4A)**. Supplementing CDM with either palmitic acid **(Fig. 4B, S6A)** or stearic acid **(Fig. 4C, S6B)** promoted growth of both strains at all concentrations ≥ 31.25 ng/ml. A 3:1 mix of palmitic and stearic acid mimicking the ratios at which they are present in BHI **(Supplementary Excel Table S1D)** promoted growth of the double mutant up to 125 ng/ml. The mix promoted growth of the wild type at all concentrations **(Fig. S6C, D)**. Unsaturated fatty acids at a concentration of 5 µg/ml inhibited growth of both strains **(Fig. S6E, F)**. Of the other saturated fatty acids (myristic, lauric, and arachidic acids), only arachidic acid promoted growth of both strains at all concentrations **(Fig. S6G, H, I)**. Collectively, our findings suggest that the *mprF* null mutant relies on exogenous palmitic acid and stearic acid for survival when *de novo* fatty acid biosynthesis is downregulated.

**Figure 4.**
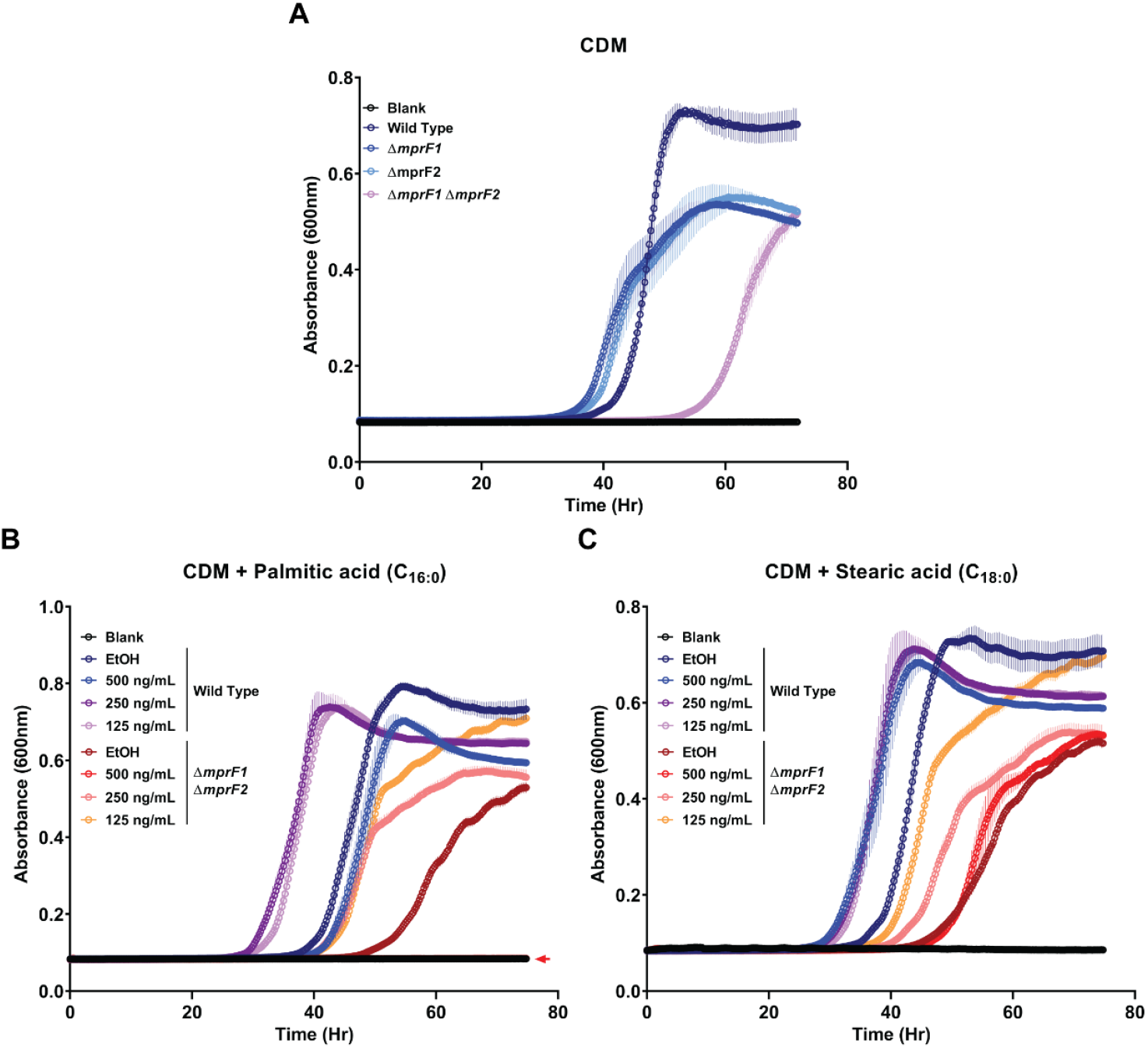
*mprF* mutants possess a growth defect when grown in chemically defined media (CDM) which can be rescued by palmitic or stearic acid. **(A)** Growth curves of wild type and *mprF* mutants grown in chemically defined media. Growth curves of wild type and ∆*mprF1* ∆*mprF2* grown under fatty acid supplementation of **(B)** palmitic acid (C_16:0_) or **(C)** stearic acid (C_18:0_). Equal volumes of ethanol (EtOH) were used as the solvent control for comparison. Each data point represents the mean ± standard error of measurement calculated from 3 biological replicates averaged from 3 technical replicates each. Δ*mprF1* Δ*mprF2* with 500 ng/mL palmitic acid in (B) is overlaid over the blank values as denoted by the colored arrow.

### L-PG depletion increases the proportion of long chain acyl-ACPs in the double mutant

*Lactococcus lactis* uses the MarR (multiple antibiotic resistance repressor) family repressor FabT to regulate expression of the *fab* gene operon (33). FabT binds to regulatory elements of the FA biosynthesis operon to repress transcription of *fab* genes in *Streptococcus pneumoniae* (34). However, in *S. pneumoniae*, FabT binds DNA only when complexed with acyl-acyl carrier protein (acyl-ACP) species that have long-chain acyl moieties (35). Acyl carrier proteins (ACPs) play essential roles in fatty acid synthesis as well as phospholipid synthesis by mediating the transfer of long fatty acyl chains from fatty acids to glycerol 3-phosphate (36, 37). *E. faecalis* FabT is 51% identical to that of *S. pneumoniae* and functions in a similar manner to *S. pneumoniae* FabT (38). Thus, we hypothesized that L-PG depletion in the double mutant leads to an accumulation of long-chain fatty acyl-ACPs, which would increase FabT affinity for the *fab* promoter, resulting in the observed suppression of *de novo* FA biosynthesis and increased dependence on exogenous fatty acids. To measure acyl-ACP levels within our strains, we took advantage of an Asp-N protease to cleave at a conserved DSLD amino acid sequence present at the acyl attachment site, leaving behind a DSL peptide connected to 4’-phosphopantetheine and an acyl-group which is then detected using mass spectrometry. The conserved nature of the Asp-N cleavage site and consistent structure of the digestion products makes quantification of acyl-ACP species possible (39, 40). *E. faecalis* contains two *acp* genes, *acpA* and *acpB*. However, only AcpA – the ACP that is involved in *de novo* fatty FA biosysthesis – possesses the conserved DSLD sequence, while AcpB – the ACP involved in uptake of exogenous fatty acids – does not (38). Thus, this method would preferentially detect acyl-AcpA species. We observed that the double mutant contains higher proportions of acyl-AcpA species containing 10 to 18 carbon atoms than the wild type with a notable decrease in acyl-AcpA species with 2 carbon atoms **(Fig. 5, S7A)**. These data confirm the hypothesis that long-chain fatty acyl-ACPs accumulate in the double mutant and provide mechanistic insight for how changes in MprF can impact global lipid homeostasis in the cell.

**Figure 5.**
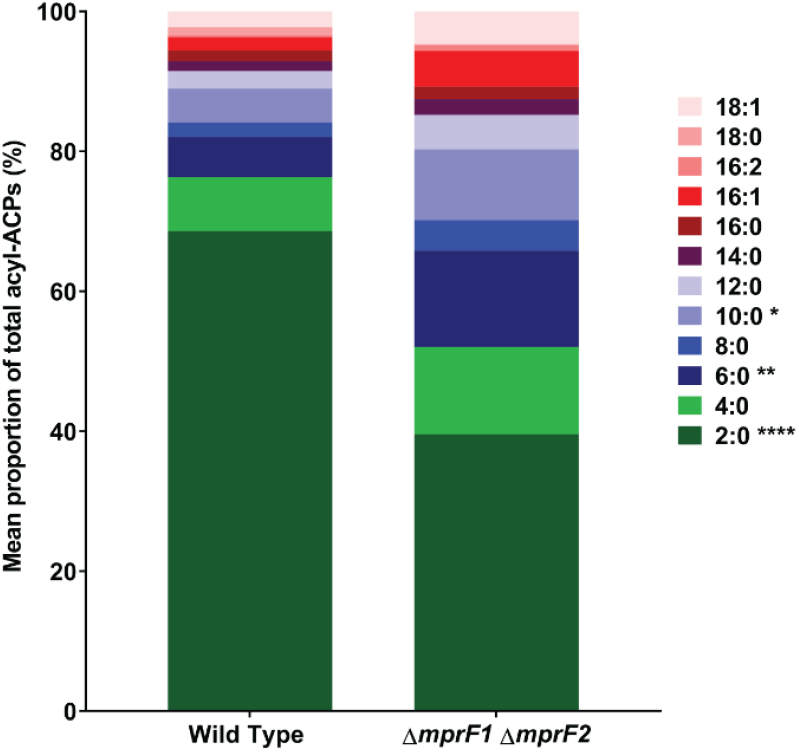
*mprF* mutants display decreased proportions of short-chain acyl-acyl carrier proteins (acyl-ACPs). Mean proportion of acyl-ACPs within WT and Δ*mprF1*Δ*mprF1* derived from normalized concentrations measured using mass spectrometry. Data from 4 biological replicates was analyzed. Values were obtained by normalizing against a 15N acyl-ACP internal standard and total of protein concentration of the respective samples. Statistical comparisons made for Δ*mprF1* Δ*mprF2* against wild type. *, p<0.05; **, p<0.001; ****, p<0.0001; Fisher’s LSD test for ANOVA. Refer to **Fig. S7A** for values of each individual species.

### Fatty acid profiles are altered in *mprF* mutants

Given the downregulation of *de novo* fatty acid biosynthesis genes in ∆*mprF1* ∆*mprF2* and the ability of exogenous fatty acids to support this mutant’s growth in CDM, it was expected that the fatty acid profile of the double mutant differs from that of the wild type in the two different media (BHI and CDM). Gas chromatography analysis of total fatty acid methyl esters (GC-FAME) was performed on lyophilized cell pellets of the wild type and ∆*mprF1* ∆*mprF2* grown in either BHI or CDM. The fatty acid profiles of the wild type and ∆*mprF1* ∆*mprF2* grown in BHI were very similar **(Fig. 6, Supplementary Excel Table S1E)**. However, when grown in CDM, ∆*mprF1* ∆*mprF2* had 11% more palmitic acid (C_16:0_), ∼11% less *cis*-vaccenic acid (C_18:1 ω7 cis_), and ∼2.5% less C_19 cyclo ω7_ acid relative to the wild type, whereas the wild type showed a decrease of 2% and 3% in palmitic acid (C_16:0_) and stearic acid (C_18:0_), respectively, and a ∼6% increase in *cis*-vaccenic acid (C_18:1 ω7 cis_) compared to growth in BHI **(Fig. 6, Supplementary Excel Table S1E)**. ∆*mprF1* ∆*mprF2* grown in CDM showed a ∼11% increase in palmitic acid (C_16:0_), a ∼7% decrease in *cis*-vaccenic acid (C_18:1 ω7 cis_), and a ∼3.5% decrease in C_19 cyclo ω7_ relative to growth in BHI **(Fig. 6, Supplementary Excel Table S1E)**. Collectively, these results suggest that the ∆*mprF1* ∆*mprF2* fatty acid composition differs from that of the wild type, in a growth medium-dependent manner. Wild type *E. faecalis* makes less C_16:0_ and more C_18:1 ω7c_ in CDM while ∆*mprF1*∆*mprF2* makes more C_16:0_ and less C_18:1 ω7c_ in CDM. These fatty acid differences could be the cause, or the effect, of the double mutant’s severely impaired growth in CDM.

**Figure 6.**
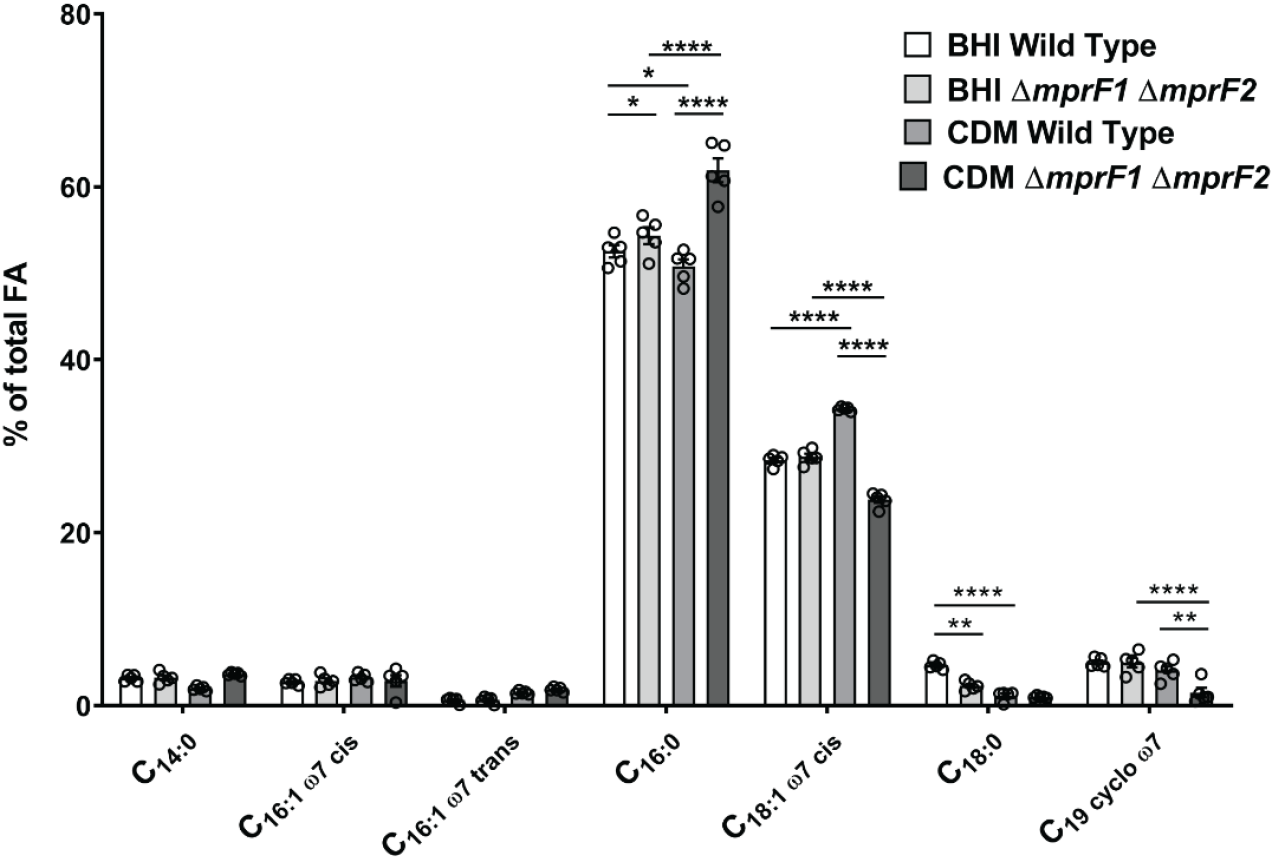
*mprF* mutants possess an altered fatty acid profile when grown in CDM as compared to the wild type. GC-FAME analysis of fatty acids in WT and ∆*mprF1* ∆*mprF2* grown in either BHI or CDM. The most abundant fatty acid species that account for at least 1% or more of the total fatty acids (FA) present within the sample are displayed here. The full list of detected fatty acid methyl esters is shown in Excel **Table S1E**. Each bar represents the mean ± standard error of measurement calculated from 5 biological replicates. *, p<0.05; **, p<0.01; ****, p<0.0001; Tukey test for ANOVA.

### *mprF* deletion has multiple functional consequences

We hypothesized that extensive lipidomic and transcriptomic alterations in the absence of *mprF* would affect cell physiology. Below we describe multiple functional changes that we observed in the *mprF* mutants.

#### Secretion

Anionic membrane phospholipids promote efficient secretion via the generalized Sec pathway (41, 42). Therefore, we hypothesized that lower PG levels in the *mprF* mutants would impair secretion. To test this hypothesis, we compared the secretion efficiency of wild type with that of the single and double *mprF* mutants. We observed reduced bulk secretion in the *mprF* mutants with the greatest decrease observed for ∆*mprF1* ∆*mprF2*, followed by ∆*mprF2* and ∆*mprF1* **(Fig. 7A)**. To validate the disruption of Sec-mediated secretion, a chimeric alkaline phosphatase was heterologously expressed in wild type and the *mprF* mutants and its secretion into the supernatant detected colorimetrically. Relative to wild type, the *mprF* mutants displayed significant impairment in secretion, with ∆*mprF1* ∆*mprF2* showing the greatest secretion defect **(Fig. 7B)**.

**Figure 7.**
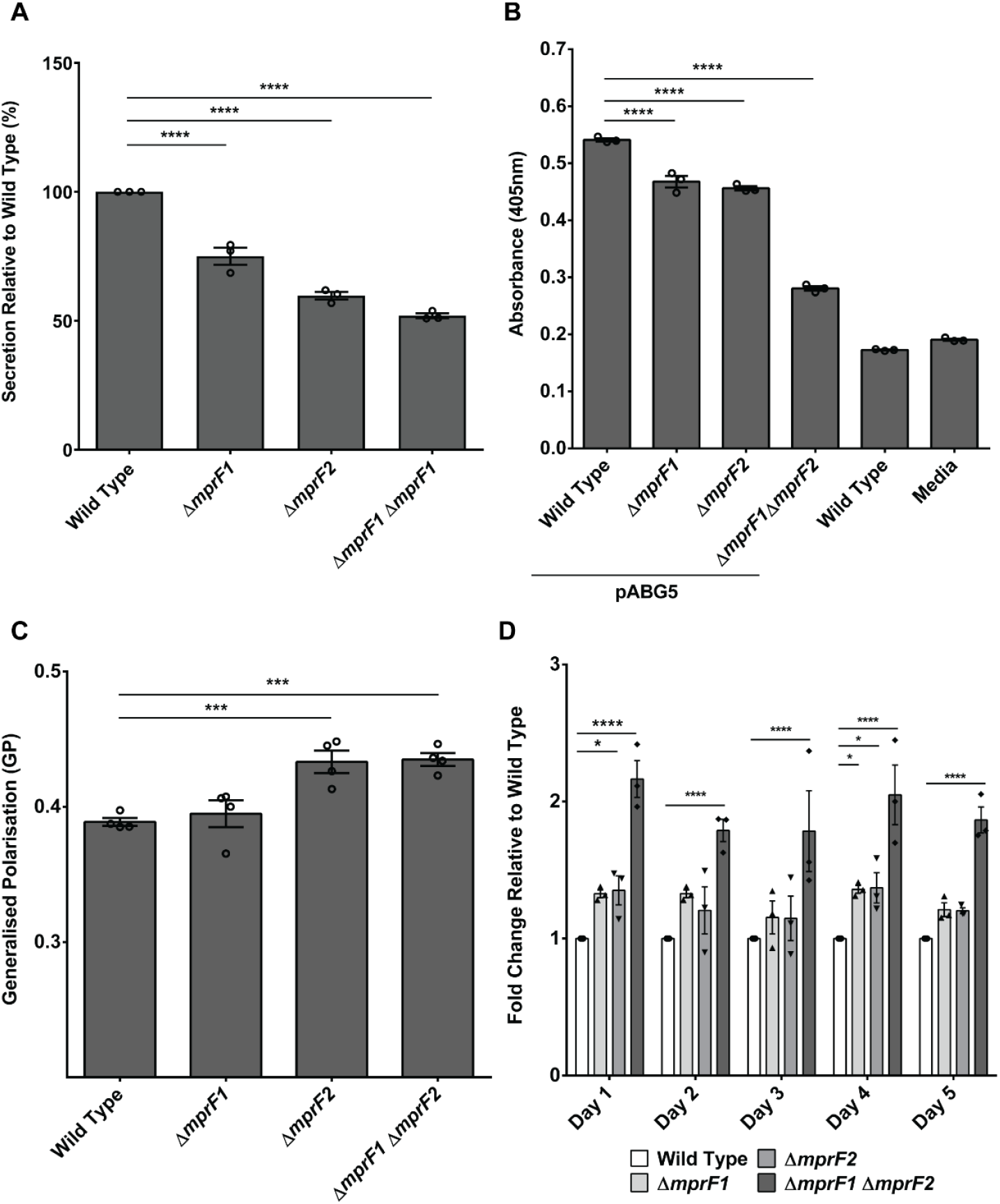
*mprF* mutants exhibit pleiotropic phenotypes. **(A)** Relative amounts of proteins secreted into the growth media by wild type and mutant strains are shown. Each bar represents the mean ± standard error of the mean calculated from 3 biological replicates averaged from 3 technical replicates each. **(B)** Relative amounts of alkaline phosphatase (PhoZF encoded on the plasmid pABG5, *E. faecalis* native PhoZ fused to the secretion domain of protein F from *S. pyogenes*) secreted into the growth media by wild type and mutant strains growth to mid-logarithmic phase are shown. PhoZF secretion was monitored by its ability to convert para-nitrophenyl phosphate (pNPP) into a colored product that can be measured by absorbance at 405 nm. Each bar represents the mean ± standard error of the mean calculated from 3 biological replicates averaged from 3 technical replicates each. **(C)** Cells were labeled with Laurdan and analyzed microscopically to assess for membrane fluidity changes. Higher GP values indicate more rigid membranes. Error bars represent the standard error of the mean of GP values from 4 separate experiments. Each experiment consists of 1 biological replicate with average GP values tabulated from >100 ROIs of cells/cell clusters. (Controls to ensure that the Laurdan assay can sensitively and accurately measure differences in membrane fluidity can be found in **Fig. S7B, C**). **(D)** Static biofilm assay with crystal violet staining across 5-days. Relative fold-change of absorbance 595 nm readings of the crystal violet stain are reported with respect to the wild type. Each bar represents the mean ± standard error of the mean calculated from 3 biological replicates averaged from 3 technical replicates each. *, p<0.05; ***, p<0.001; ****, p<0.0001. All data sets were analyzed using Fisher’s LSD test for ANOVA.

#### Membrane fluidity

The fluidity of the cell membrane is influenced by the amount of unsaturated lipids present, where a higher degree of unsaturation results in more fluid membranes (43). Based on the reduction of unsaturated L-PG in the *mprF* mutants **(Fig. 2, S1C)** and alterations in other lipid classes, we hypothesized that membrane fluidity might be affected as well. Membrane fluidity was assayed using a fluorescent dye sensitive to fluidity changes, Laurdan. Laurdan inserts into membrane bilayers and, depending on how liquid ordered (L_o_) or disordered (L_d_) the local lipid environment is, its emission spectra will blue-shift (in L_o_ regions) or red-shift (in L_d_ regions) respectively (44, 45). This spectral shift can be detected by measuring the green and blue wavelengths and expressing the readings as a ratio, generalized polarization (GP), where higher GP values imply more rigid membranes and lower GP values imply more fluid membranes. By staining late stationary phase cultures of the wild type and *mprF* mutants with Laurdan and imaging cells by microscopy, we observed that ∆*mprF2* and ∆*mprF1* ∆*mprF2* had higher GP values, indicating slightly more rigid membranes in these mutants compared to the wild type, while ∆*mprF1* had no significant difference in GP values **(Fig. 7C)**. Hence, the lipidomic changes observed in ∆*mprF1* ∆*mprF2* are correlated with lower membrane fluidity.

#### Biofilm formation

In *E. faecalis* strain 12030, the loss of *mprF* enhances biofilm formation (14). To determine whether loss of *mprF* has the same effect in strain OG1RF, we used crystal violet staining to assay biofilm formation from wild type and *mprF* mutants grown in microtiter plates. The stain detects adherent biomass. We observed slight increases in biofilm formation for the ∆*mprF1* and ∆*mprF2* mutants, with the increase being significant for ∆*mprF1* ∆*mprF2* (P ≤ 0.001) across all daily timepoints over a 5-day period **(Fig. 7D)**.

## Discussion

The major conclusion of this study is that depletion of a cationic membrane phospholipid (L-PG) leads to unexpected lipidomic, transcriptomic, and functional changes in *E. faecalis*, and that the two MprF paralogs of *E. faecalis* contribute differently to the observed changes. Specifically, MprF2 is not only more important for protection against CAMP-mediated killing than MprF1, but is also involved in global lipid homeostasis and cell function.

In the current study, we used previously established mass spectrometry-based methods (10) to analyze various lipids in *E. faecalis* OG1RF, ∆*mprF1*, ∆*mprF2*, and ∆*mprF1* ∆*mprF2*. One advantage of such methods is that they are much more sensitive than traditional thin layer chromatography (TLC), enabling a more comprehensive understanding of lipid homeostatic shifts that are dependent upon *mprF* (14). When we compared the lipid compositions of *mprF* deletion mutants with the parent OG1RF strain, we found that, as expected, deletion of *mprF2* resulted in a complete absence of L-PG from the membrane. The observed absence of L-PG in ∆*mprF2* is consistent with our previous work showing that ∆*mprF2* is significantly more sensitive to killing by human β-defensin 2 (hBD2) than ∆*mprF1* (9). Unlike previous reports, however, we found that *mprF1* also contributes to L-PG production in *E. faecalis*, a key phenotypic difference that motivated us to explore changes elsewhere in the *E. faecalis* lipidome. While we observed that L-PG is only slightly reduced in ∆*mprF1* (*mprF2* present), L-PG is completely absent in ∆*mprF2* (*mprF1* present). The unexpected observation that MprF1 alone does not synthesize L-PG suggests that MprF2 expression is necessary for MprF1 to function. This possibility warrants further investigation.

To our surprise, we also discovered that ∆*mprF2 and* ∆*mprF1*∆*mprF2* have significantly lower PG levels than wild type and ∆*mprF1*. This is a novel and unexpected finding because *mprF* acts downstream of *pgsA*, one of two genes (along with *pgpA*) involved in PG synthesis **(Fig. 8)**.

**FIgure 8.**
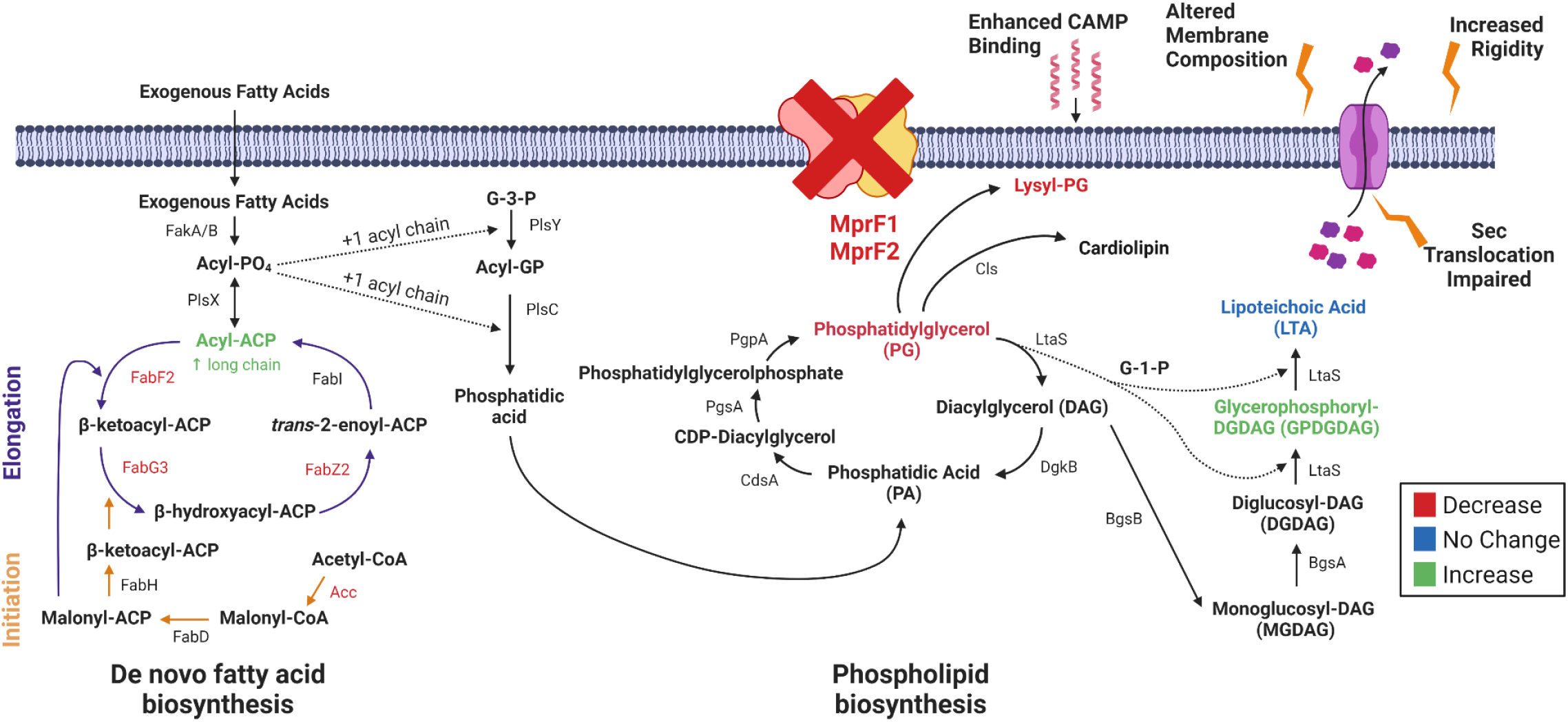
A working model for lipidomic, transcriptomic, and phenotypic consequences following *mprF* deletion. The loss of *mprF* leads to wnregulation of fatty acid biosynthesis genes and decreases in lysyl-PG (L-PG) and PG coupled with increases in GPDGDAG and a phosphorus-ntaining lipid of unknown identity. These large lipidomic changes lead to functional impairment in Sec-mediated secretion, increased membrane idity, and enhanced CAMP binding. Created with BioRender.com.

Concurrently, levels of GPDGDAG, D-ala-GPDGDAG, and a phosphorous-containing lipid increase, possibly a cellular response to compensate for the loss of PG and L-PG and maintain cell membrane integrity. DGDAG is a membrane lipid that serves as the lipid anchor of lipoteichoic acids (LTA) in Gram-positive bacteria. As the phosphoglycerol headgroup is consumed to form polymeric LTA, DAG accumulates in the membrane (30). It has been suggested that the consumption of phosphorous could be offset by an accumulation of DGDAG in *Staphylococcus haemolyticus* and *Staphylococcus epidermidis* (46). GPDGDAG is a glycolipid formed as an intermediate during synthesis of lipoteichoic acid (LTA) in *S. aureus* (47). A lipidomic comparison between daptomycin-resistant *E. faecalis* strain R712 and daptomycin-sensitive strain S613 revealed that R712 had less PG, L-PG, and CL than S613 but more MGDG and DGDG, and that this glycolipid upregulation is consistent with an increase in GPDGDAGs reported earlier by another group (20, 48). Based on these earlier findings and our own results, we conclude that these phospholipids and glycolipids are linked by common pathways in *E. faecalis*. Additional experiments are needed to determine the identity of the phosphorus-containing lipid, and to fully understand the roles of it, along with DGDAG and GPDGDAG, in *E. faecalis* membrane homeostasis and function.

We show that long-chain acyl-ACPs, precursors for phospholipid synthesis, accumulate in the absence of MprF, which we speculate is a consequence of the loss of major lipids such as PG and L-PG. These long-chain acyl-ACPs activate the transcriptional repressor FabT, which represses *de novo* fatty acid biosynthesis, resulting in increased dependence on exogenous fatty acids for growth. In *E. faecalis*, acyl-AcpA was recently shown to enhance FabT binding to DNA (49). We thus propose a model in which deletion of *mprF1* and *mprF2* mediates lipidomic and transcriptomic changes via enhanced FabT activity resulting from an accumulation of long-chain acyl-AcpA species **(Fig. 8)**.

As a consequence of these lipidomic and transcriptomic changes, the cells manifest a secretion defect, increased membrane rigidity, and enhanced CAMP binding. The secretion machinery of *B. subtilis* requires interfacial regions of high and low fluidity to function optimally (44). As membrane fluidity influences secretion, the impaired secretion that we observed in ∆*mprF1*, ∆*mprF2, and* ∆*mprF1*∆*mprF2* is consistent with findings in *B. subtilis* as well as with earlier studies demonstrating that efficient secretion requires anionic phospholipids (41, 42, 50, 51). Another study which investigated factors governing *Listeria monocytogenes* secretion of listeriolysin demonstrated a secretion defect in an *mprF* mutant, further implicating MprF in bacterial secretion (52). Collectively, these observations support a model of efficient secretion requiring both modified anionic phospholipids and optimal membrane fluidity in *E. faecalis*.

We previously reported that all L-PG species in laboratory-grown *E. faecalis* are unsaturated, i.e. they contain at least one carbon-carbon double bond in a constituent acyl chain (24). Here we show that all L-PG species are absent from ∆*mprF2* and ∆*mprF1*∆*mprF2*. As acyl chain unsaturation is correlated with membrane fluidity (53), our results suggest that the lower membrane fluidities of these two mutant strains are due to the lack of unsaturated L-PG species. Although the ratio of saturated-to-unsaturated acyl chains is similar for the wild type and the double mutant, a recent study suggests that acyl chain remodeling in different phospholipid classes affects membrane properties differently (22). In that study, out of 27 lipid species identified in *Methylobacterium extorquens*, as few as 8 were highly variable over all of the conditions tested (various temperatures, osmotic and detergent stresses, carbon sources, and cell densities). Thus, only a fraction of the lipidome was involved in adaptive remodeling. Different sets of lipidomic features (phospholipid class, acyl chain saturation, and acyl chain length) were involved in responses to changes in temperature, high salt concentrations, and stationary growth phase. One of the more striking observations was that varying the degree of acyl chain saturation in PE, PG, or phosphatidylcholine (PC) had varying effects on lipid packing, a property that is correlated with other physical properties including viscosity (54). Cells maintain optimal membrane properties when exposed to environmental challenges (e.g. changes in temperature, acidic pH, osmotic stress) through an adaptive response known as homeoviscous adaptation, which involves changes to the acyl chain composition of membrane lipids (55). Consistent with the *M. extorquens* study, our findings support the concept that lipid class-dependent acyl chain remodeling provides a mechanism by which to optimize the cell membrane’s biophysical properties, thus greatly improving our understanding of membrane adaptation.

In summary, our findings therefore suggest roles for MprF beyond mediating resistance to cationic antimicrobials via L-PG synthesis. Our study underscores the need to consider broader consequences of mutations involving lipid-related genes, as they may lead to unanticipated changes within other lipid classes as part of an adaptive response involving global lipid remodeling. Such adaptations need to be considered when devising novel antimicrobial strategies that target membrane lipids. The lipidome remodeling we report here also underscores the importance of mass spectrometry-based lipidomics in understanding the behavior of one of the most clinically significant opportunistic pathogens.

## Materials and Methods

Strains, growth conditions, RT-qPCR, growth kinetics, live/dead staining, RNA sequencing, cloning methods and methods for MS analysis of the TLC spots are detailed in the **Supplementary Text 1B**.

### Analysis of membrane lipid content

Lipids were extracted from lyophilized cell pellets from late stationary phase cultures using a modified Bligh & Dyer method in which the extraction solvent contained chloroform/methanol in a ratio of 1:2 (v/v) as previously described (10). For method validation and quantification, known amounts of internal standards for phosphatidylglycerol (PG) and lysyl-PG (L-PG) were added to the samples. Nine hundred microliters of chilled extraction solvent containing internal standards (Avanti polar lipids, Alabaster, AL, USA) was added to the dry cell pellets except monoglucosyl-diacylglycerol (MGDAG) 34:1 which was used as a surrogate external standard for diglucosyl-diacylglycerol (DGDAG) instead **(Table S1B)**.

Lipid extraction was then carried out as previously described (10). The dried lipid extract was resuspended in a mixture of chloroform and methanol (1:1 v/v), to a final lipid concentration of 10 mg/ml. This solution was stored at -80 °C until the mass spectrometry analysis was performed. PG and L-PG in *E. faecalis* were quantified by LC-MS/MS using multiple reaction monitoring (MRM) using a previously described methodology (10). An Agilent 6490 QqQ mass spectrometer connected to a 1290 series chromatographic system was used with a Kinetex^®^ 2.6 μm HILIC column (100 Å, 150 × 2.1 mm) (Phenomenex, USA). Electrospray ionization (ESI) was used to ionize lipids. Each lipid molecular species was analyzed using a targeted multiple reaction monitoring (MRM) approach containing transitions for known precursor/product mass-to-charge ratio (m1/m3). Signal intensities were normalized to the spiked internal standards (PG 14:0 and L-PG 16:0) to obtain relative measurements and further normalized against the initial dry cell pellet weight, as described previously (10).

To determine the species of DGDAG present in *E. faecalis*, untargeted analysis of lipid extracts of WT, ∆*mprF1*, ∆*mprF2* and ∆*mprF1* ∆*mprF2* was carried out. Lipid extracts were analyzed using an Agilent 6550 QToF mass spectrometer connected to a 1290 series chromatographic system with a Kinetex^®^ 2.6 μm HILIC column (100 Å, 150 × 2.1 mm) (Phenomenex, USA). The QToF instrument was set to positive ion mode, at an electrospray voltage of -3500 V (Vcap), a temperature of 200 °C, a drying gas rate of 14 L/min. Spectra were acquired in auto-MS2 mode with MS1 acquisition rate at 4 spectra/s and the MS2 acquisition rate at 20 spectra/s with fixed collision energy at 40 eV. The list of detected species of DGDAG in the samples can be found in the **Supplementary Excel Table S1A**.

Due to the absence of suitable internal standards, semi-quantitative analysis of DGDAG was carried out instead. Lipid extraction was performed as described above without addition of internal standards. Analysis of DGDAG lipid species was performed by LC-MS/MS via MRMs using monoglucosyl-diacylglycerol (MGDAG) 34:1 as a surrogate standard **(Table S1B)** for external calibration curves. Measurements of MGDAG 34:1 dilution from 0.2 ng/mL to 1000 ng/mL were used to construct external calibration curves to estimate the levels of DGDAG. Estimated DGDAG levels were then normalized against dry cell pellet weight of the respective samples. The MRM transitions for DGDAG molecular species and MGDAG 34:1 are listed in **Supplementary Excel Table S1B**. The mobile phase gradients used for all experiments are as previously described (10). For semiquantitative analysis of glycerophosphoryl-diglucosyl-diacylglycerol (GPDGDAG), lipid extracts were analyzed using an Agilent 6550 QToF mass spectrometer connected to a 1290 series chromatographic system with a Kinetex^®^ 2.6 u HILIC column (100 Å, 150 × 2.1 mm) in negative mode. The most abundant GPDGDAG was quantified with the following transition (MS1 m/z: 1071.6, MS2 m/z: 153.0). Integrated peak areas were normalized against cell weight.

### RNA Sequencing

Sequencing of RNA was done from OG1RF, ∆*mprF1*, ∆*mprF2*, and ∆*mprf1*∆*mprF2* strains. Detailed methods are described in the supplementary information file. RNAseq files are available on NCBI, Sequence Read Archive (SRA) (Accession: PRJNA634972).

### Analysis of acyl-ACP content

Quantification of acyl-ACPs were done as previously described with the following modifications (39, 40). 20 mL of overnight cultures were pelleted, treated with 10 mg/mL of lysozyme solution for 1 hour and lysed using a probe sonicator at 40% amplitude for 2 minutes (pulsed at 30s on and 30 s off) for 2 cycles. Unlysed cells were removed by centrifugation at 15 700 rcf for 30 minutes at 4_o_C and the concentration of proteins within the clarified lysate was then measured using the Qubit Protein Assay (Thermofisher Scientific, USA) according to the manufacturer’s instructions. 50 µL of _15_N acyl-ACP standards (Holo, 2:0, 3:0, 4:0, 6:0, 8:0, 10:0, 12:0, 13:0, 14:0, 16:0, 18:0, 18:1) were then spiked into the sample at 5 µM equimolar concentrations. Proteins were precipitated from standard spiked lysates via TCA-precipitation and resuspended in 50mM MOPS buffer, pH 7.5. Resuspended proteins were then treated with the ASP-N protease at 20:1 ratio (protein:enzyme) at 37_o_C overnight. Reaction was then stopped by addition of methanol to a final concentration of 50%. These samples were then analysed using LC-MS with MRMs using previously described solvent gradients and MRM parameters (39). An Agilent 6495A QqQ mass spectrometer connected to a 1290 series chromatographic system was used with a Discovery BIO Wide pore C18-3 (10cm x 2.1mm, 3uM particle size) (Supelco, USA). Electrospray ionization (ESI) was used to ionize lipids. Acquisition was carried out with the following source parameters: gas temperature: 290_o_C, gas flow: 12 L/min, nebulizer: 30 psi, sheath gas heater: 400_o_C, sheath gas flow: 11 L/min, capillary: 4500 V, Vcharging: 1500 V. Signal intensities were normalized to the spiked internal standards to obtain relative measurements and further normalized against the initial protein concentration.

### Fatty acid methyl esters (FAME) analysis

Late stationary phase cultures of wild type and ∆*mprF1*∆*mprF2* grown in either BHI (overnight) or CDM (72 hours) were lyophilized and sent together with powdered BHI for GC-FAME analysis at the Identification Service of Deutsche Sammlung von Mikroorganismen und Zellkulturen GmbH (DSMZ), Braunschweig, Germany. Cellular fatty acids were converted into fatty acid methyl esters (FAME) and analyzed by GC-MS using used C_21:0_ FAME in a defined amount per biomass as internal standard for normalization.

### Radiolabeling and thin layer chromatography (TLC)

Radiolabeling of lipids were performed as previously described with the following modifications (56) [_14_C]-acetate or [_32_P]-disodium phosphate (Perkin Elmer, USA) was added into 5 mL of media at 0.2 µCi/mL or 1 µCi/mL respectively before culturing strains overnight at 37_o_C for 16-18 hours at static conditions. Lipids were then extracted as previously described and resuspended in 50 μL of chloroform-methanol solution (1:1 v/v) (10). 10 μL of lipid extracts were mixed with 2 mL of Ultima Gold™ scintillation fluid (Perkin Elmer, USA), and radioactive counts were measured using a MicroBeta2^®^ scintillation counter (Perkin Elmer, USA). The lipid extracts were spotted on to silica-gel coated TLC plates (Merck, USA) and normalized according to the scintillation counts. TLC plates were developed in pre-equilibrated TLC chambers with chloroform:methanol:water (65:25:4) solvent system for 1-dimension (1D) TLCs. For 2-dimensional (2D) TLCs, TLC plates were developed using chloroform:methanol:water (65:25:4) solvent system for the first dimension and chloroform:hexane:methanol:acetic acid (50:30:10:5) for the second dimension. TLC plates were then visualized by exposure to a storage phosphor screen (GE healthcare, USA) overnight, and read using a Storm Phosphorimager (GE healthcare, USA). For iodine and ninhydrin-stained TLC plates, no radiolabeling was carried out and TLC spots were normalized based on dry cell weight instead. Iodine crystals (Sigma-Aldrich, USA) were used to develop TLC plates in a chamber, while for ninhydrin staining, ninhydrin was applied to the plates, allowed to air dry before heating with a hairdryer till spots appeared.

### SDS-PAGE and western blot

SDS-PAGE and western blot were performed as described in a previous study (57). 4-12% or 12% NuPAGE^®^ Bis-Tris mini gel in a XCell SureLock^®^ Mini-Cell filled with either 1x MES or 1x MOPs SDS running buffer (Invitrogen, USA) was used and run at 140 V for 90 min. Proteins were transferred to nitrocellulose membranes using the iBlot_TM_ Dry Blotting System (Invitrogen, USA) according to the manufacturer’s protocol. The antibodies and developing solutions used are shown in **Table S1E**.

### Bulk secretion assay

Late stationary phase cultures were prepared by growing cultures in BHI broth overnight for 16-18 hours at 37°C, static conditions. OD_600_ readings of the cultures were measured. Cell-free supernatants were obtained by centrifugation at 6,000 rcf for 5 minutes at 4_o_C and filtering the supernatants into fresh Eppendorf tubes using 0.2 μm syringe filters. 1.6 mL of filtered supernatant was mixed with 400 μL of 100% w/v tricholoroacetic acid (TCA) solution (1:4 ratio of TCA to sample) and incubated at 4_o_C for 10 minutes. Tubes were centrifuged at 20,000 rcf for 15 minutes at 4_o_C. The precipitated protein pellet was washed once with 2 mL of 100% ice-cold acetone and placed on a 98_o_C heat block to evaporate residual acetone. The pellets were resuspended in 500 μL of PBS. Twenty-five microliters of these protein solutions were used for estimation of protein content using the Pierce BCA Protein Assay Kit (Thermoscientific, USA) in a microtiter plate format according to the manufacturer’s protocol. Protein concentrations of the samples were then normalized to OD_600_ of 1.0 based on the respective OD_600_ readings of the individual cultures.

### Alkaline phosphatase (AP) secretion assay

Secretion of the strains were monitored by its ability to secrete a chimeric alkaline phosphatase, PhoZF (*E. faecalis* native PhoZ fused to the secretion domain of protein F from *S. pyogenes*) PhoZF secretion monitored by its ability to convert para-nitrophenyl phosphate (pNPP) into a colored product that can be measured by absorbance at 405 nm. Mid-log phase cultures of strains harboring the pABG5 plasmid containing the chimeric alkaline phosphatase enzyme (PhoZF) were normalized to OD_600_ of 0.5. Cell-free supernatants were obtained by centrifuging samples at 6,000 rcf for 5 minutes at 4_o_C and filtering the supernatant through 0.2 μm syringe filters. 25 μL of supernatant was added to 200 μL of 1M Tris-HCl, pH 8.0, in a 96-well microtitre plates. 25 μL of 4 mg/mL para-nitrophenyl phosphate (pNPP) (Sigma-Aldrich, USA) was then added to each well to start the reaction. The plate was then placed into a Tecan Infinite_©_ M200 Pro spectrophotometer and incubated at 37_o_C with the absorbance read at 405 nm every 10 minutes for 18 hours.

### Analyzing membrane fluidity by Laurdan staining (microscopy)

Late stationary and mid-log phase cultures were normalized to OD_600_ of 0.7 in PBS and incubated with 100 μM of Laurdan for 10 minutes at 37_o_C. Cells were washed twice with PBS and 10 μL of the cell suspension was spotted onto PBS-agarose pads (1% w/v) mounted on glass slides. Coverslips were placed over the agarose pads and sealed using paraffin wax.

Slides were imaged using a Zeiss LSM 880 Laser Scanning Microscope with Airyscan, using a Plan-Apochromat 63x/1.4 Oil DIC objective with an incubation chamber set to 37_o_C. The slides were equilibrated for 10 minutes within the chamber before imaging and excited using a 405nm laser with emission collected between 419-455nm (blue) and 480-520nm (green) simultaneously. Digital images were acquired using the Zen (Zeiss) software and analyzed using ImageJ. Using ImageJ, regions of interests (ROIs) of individual cells or cell clusters were selected and mean fluorescence intensities (MFIs) of each ROI for each channel were measured and tabulated in Microsoft excel. Using the following formula, the average GP values for each ROI were calculated and then plotted using Graphpad Prism software:

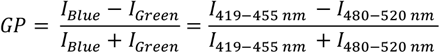

Laurdan was validated to be responsive to changes in membrane fluidity via control experiments subjecting stained cells to a gradient of temperatures, and membrane fluidizer, benzyl alcohol **(Fig. S7B, C)**.

### Daptomycin minimum inhibitory concentration (MIC)

Mid-log phase cultures of the strains were tested for their daptomycin minimum inhibitory concentration (MIC) using the microplate broth dilution methods as previously described (10).

### Static biofilm assay with crystal violet staining

Static biofilm assays with crystal violet staining were performed as previously described on the wild type, ∆*mprF1*, ∆*mprF2*, and ∆*mprF1* ∆*mprF2* across a 5-day period (58).

### Antimicrobial Peptide (AMP) Susceptibility Assays

Overnight OG1RF cultures were subcultured in BHI liquid broth at a 1:10 dilution and grown to mid-log phase (OD_600_ 0.5±0.05). These bacterial cultures were then normalized to OD_600_ 0.5 and harvested by centrifugation at 6,000 rcf for 5 minutes at 4_o_C. The supernatant was discarded and the remaining cell pellet was washed and resuspended in 1 mL of 0.01 M low-salt phosphate buffer (PB). Bacterial resuspensions were then serially diluted 500-fold and 25 μL of each sample was added twice into a 96-well microtiter plate, for a total of 2 technical replicates. An equal volume of human β-defensin 2 (hBD2) or LL-37 (Peptide Institute Inc., Japan) was added to the samples and incubated statically for 2 hours at 37_o_C **(Table S1G)**. Serial dilution was then performed up to 10_-8_ using 1X sterile phosphate buffered saline (PBS). 5 μL of bacterial suspension from each well was then spotted 3 times onto a BHI agar plate, for a total of 6 technical replicates per sample. The spotted agar plates were then incubated statically overnight at 37_o_C and surviving bacteria were determined by CFU enumeration.

## Supporting information

Supplementary Text 1A-B, Supplementary References

## Acknowledgments

This work was supported by the National Research Foundation and Ministry of Education (MOE) Singapore under its Research Centre of Excellence Program, and by an MOE AcRF Tier 1 grant (MOE2017-T1-001-269) and a National Medical Research Council (NMRC) grant (OFIRG20nov-0079), both awarded to KAK. Work in the MRW laboratory was supported by grants from the National University of Singapore via the Life Sciences Institute (LSI), the National Research Foundation (NRFSBP-P4) and the NRF and A*STAR IAF-ICP I1901E0040. SLC was supported by National Medical Research Council (NMRC) grants (NMRC/CIRG/1358/2013 and NMRC/OFIRG/0009/2016). SSC acknowledges support from the Singapore Ministry of Health National Medical Research Council under its Open Fund Individual Research Grant (MOH-000145). DKA and SAM acknowledge support from U.S. Department of Agriculture-Agricultural Research Service and the National Science Foundation Major Research Instrumentation program, award #DBI-1427621 that funded a mass spectrometer used in aspects of the project. We thank Adeline Mei Hui Yong for help with RNA extractions and Tan Wee Boon for assisting with the radiolabeling of lipids.

## Graphical Abstract

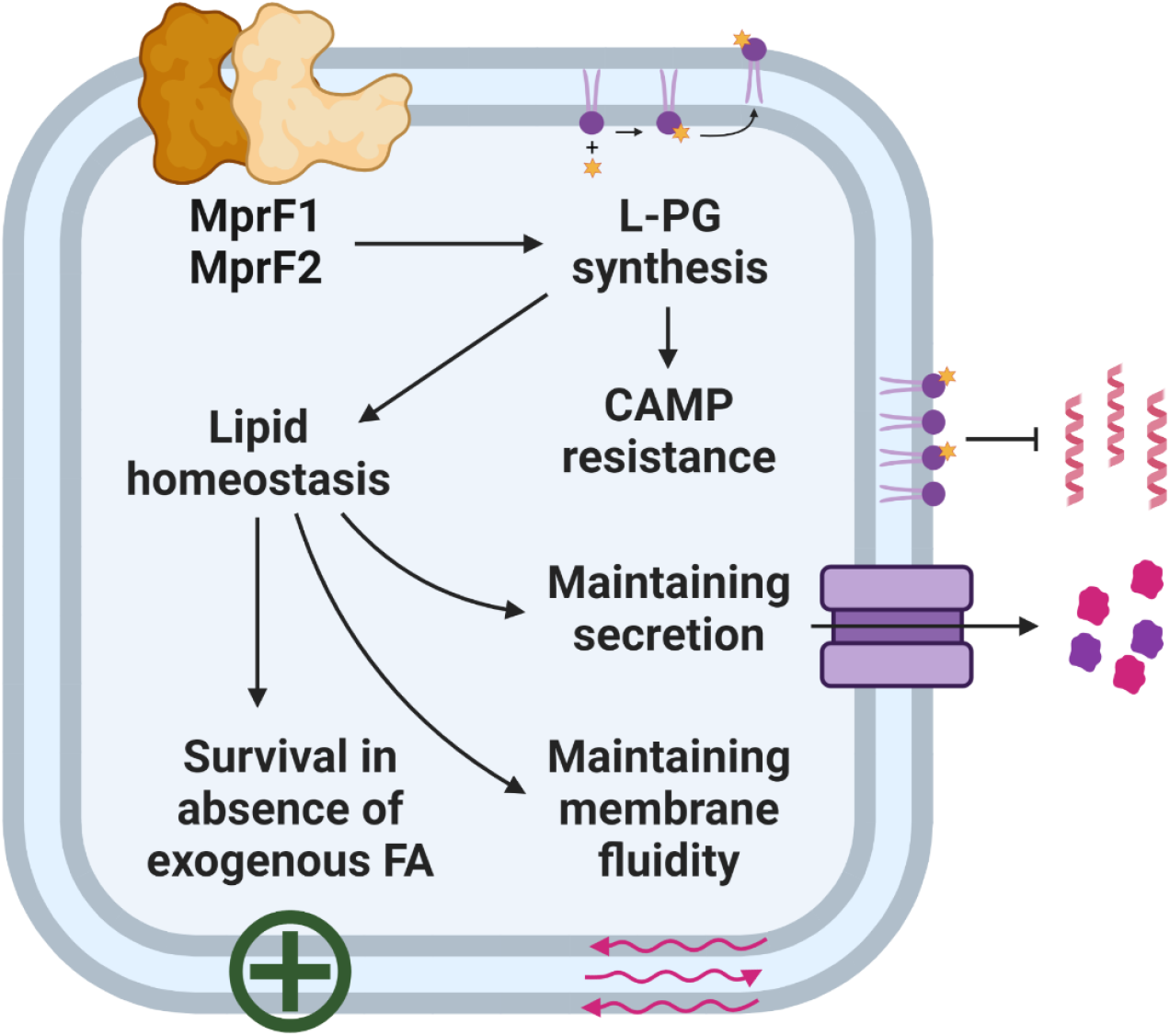

## Notes

### Competing Interest Statement

The authors have declared no competing interest.

