## Supplementary Figures S1-S7 for "Depleting cationic lipids involved in antimicrobial resistance drives adaptive lipid remodeling in *Enterococcus faecalis*"

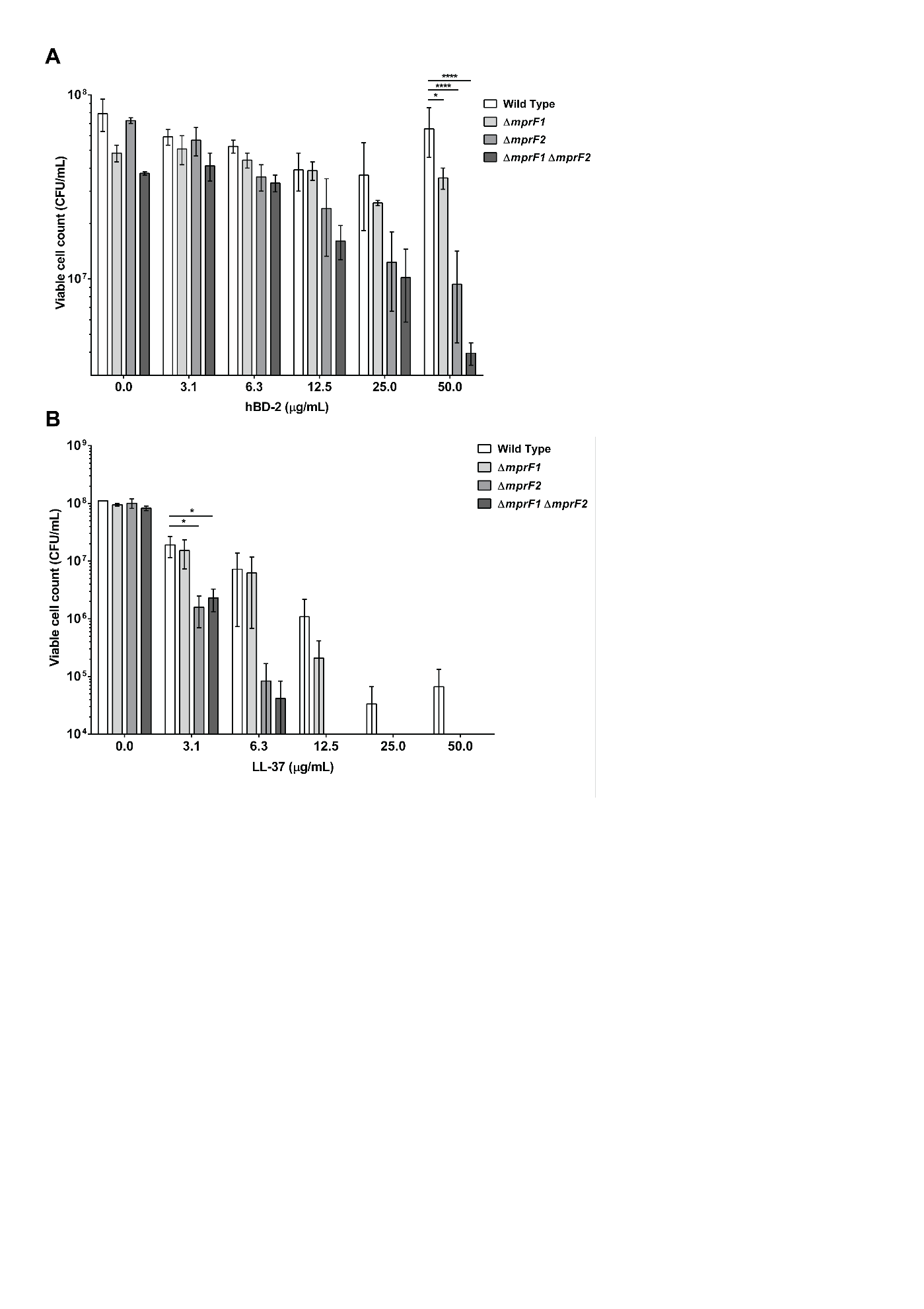

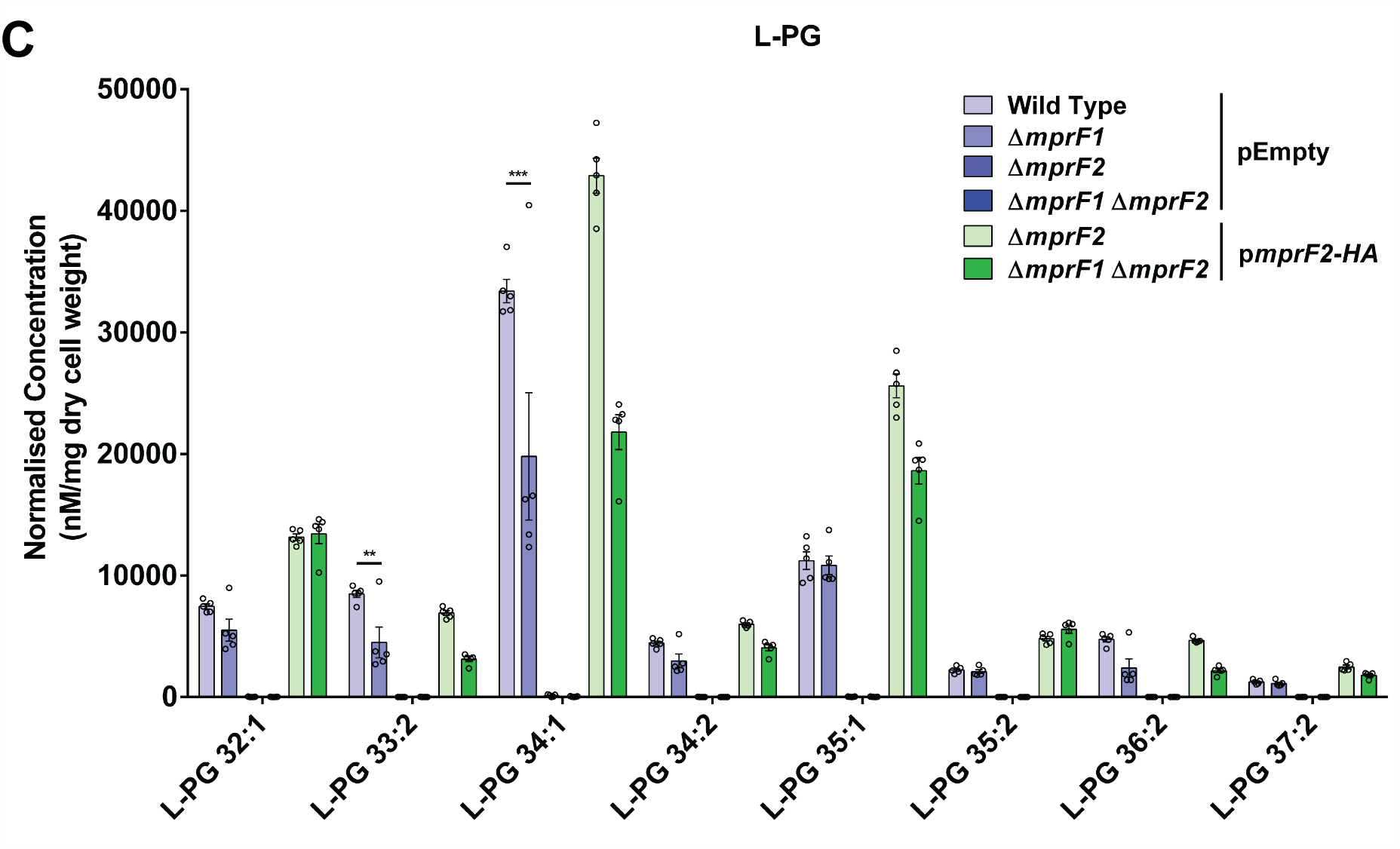

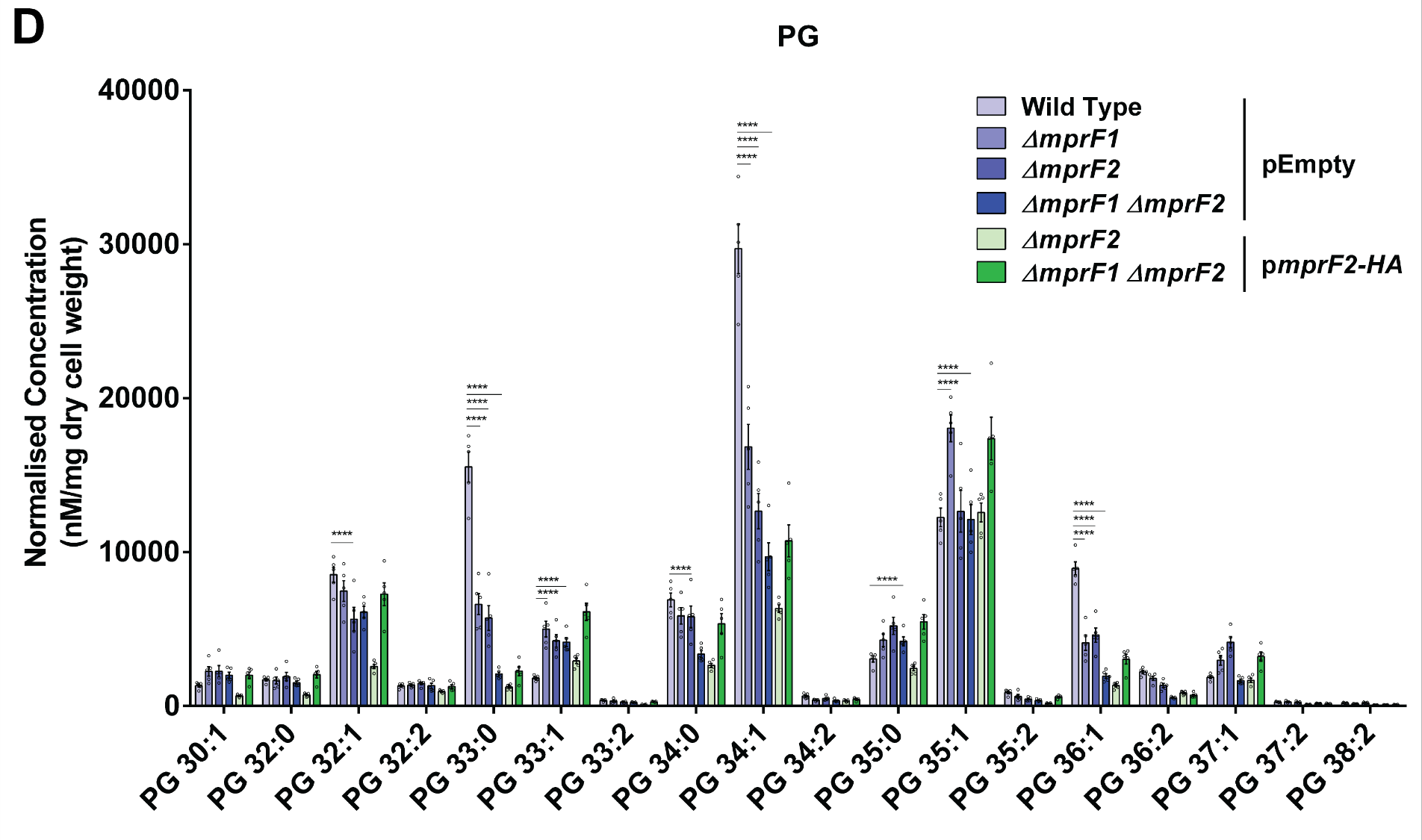

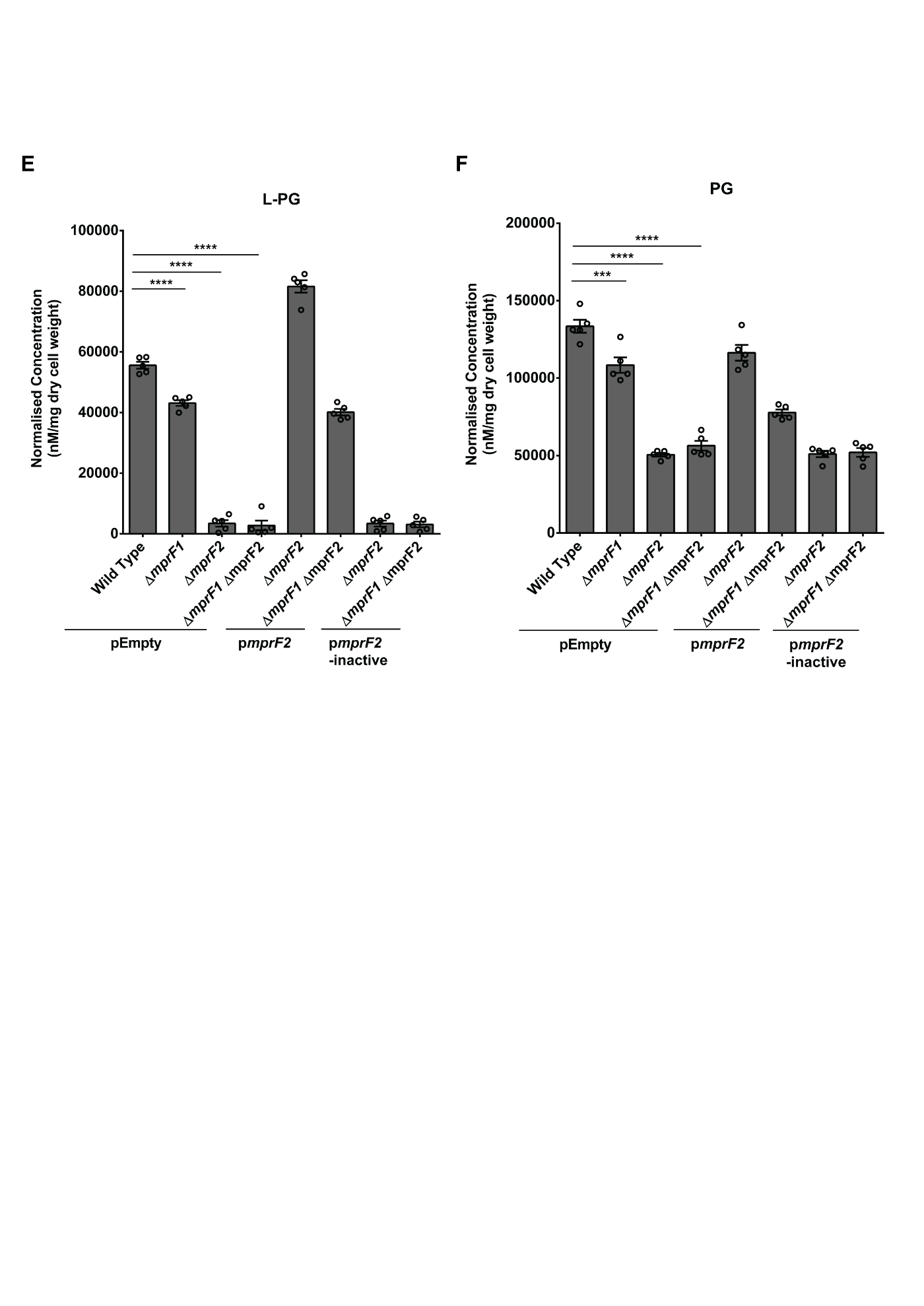

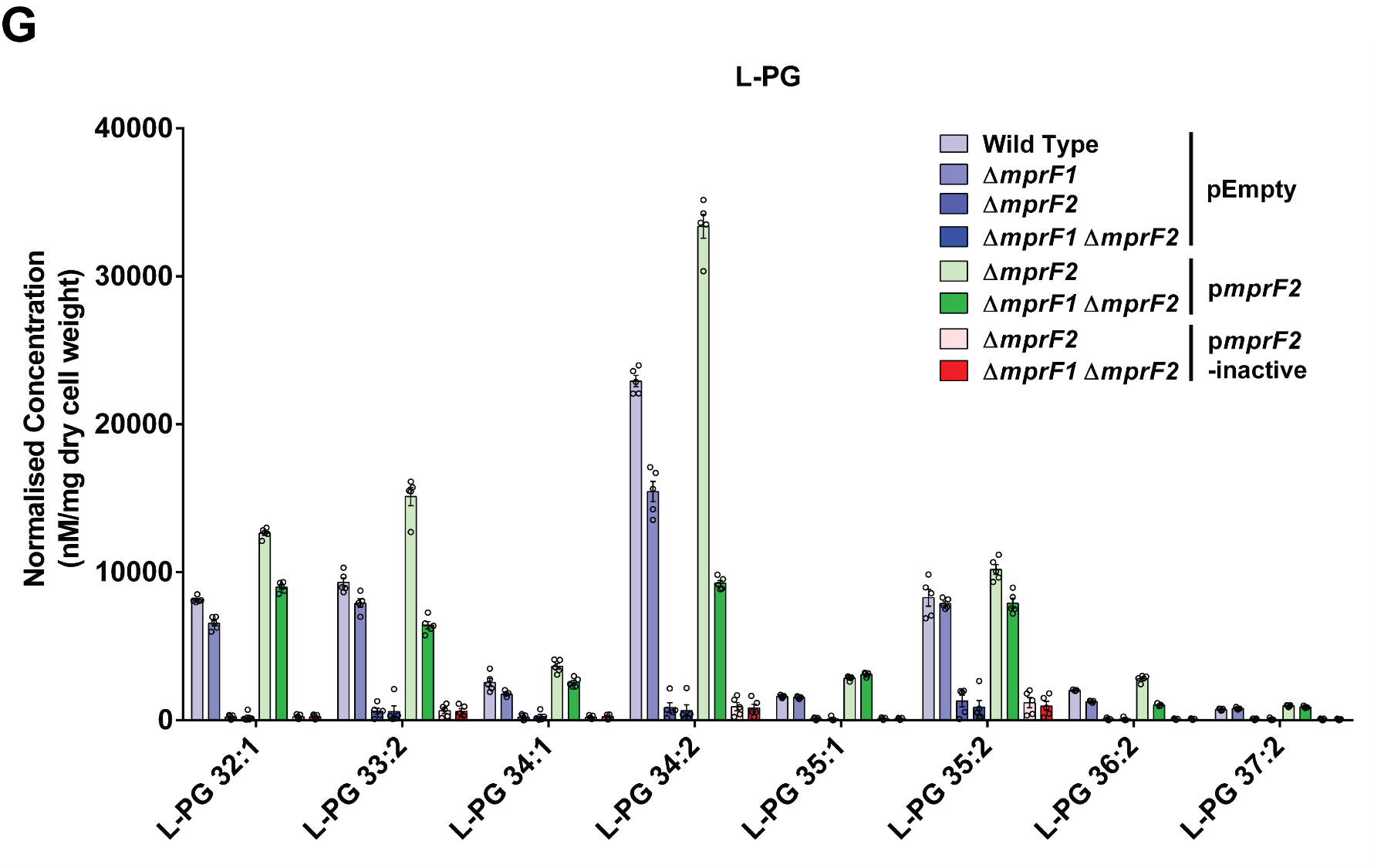

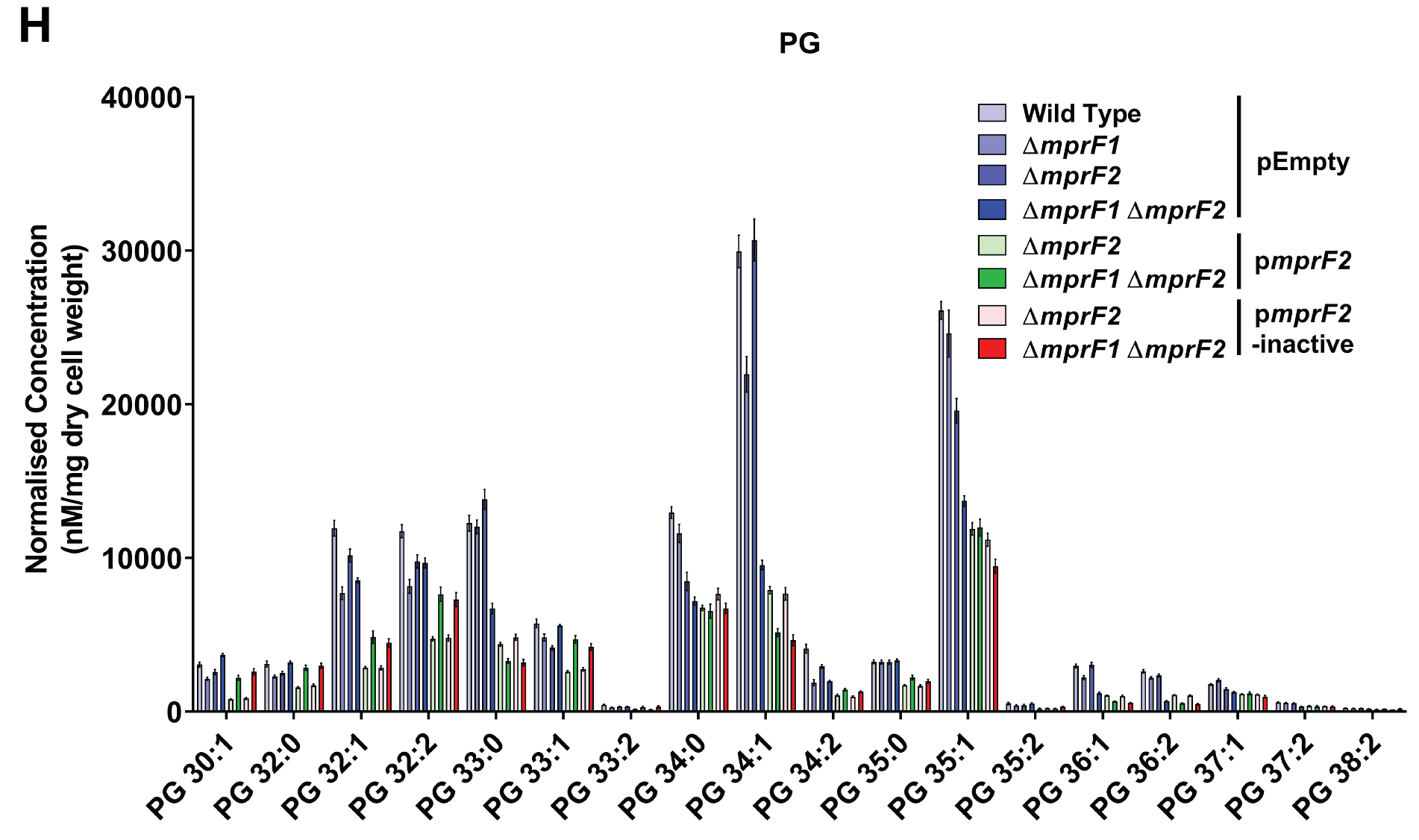

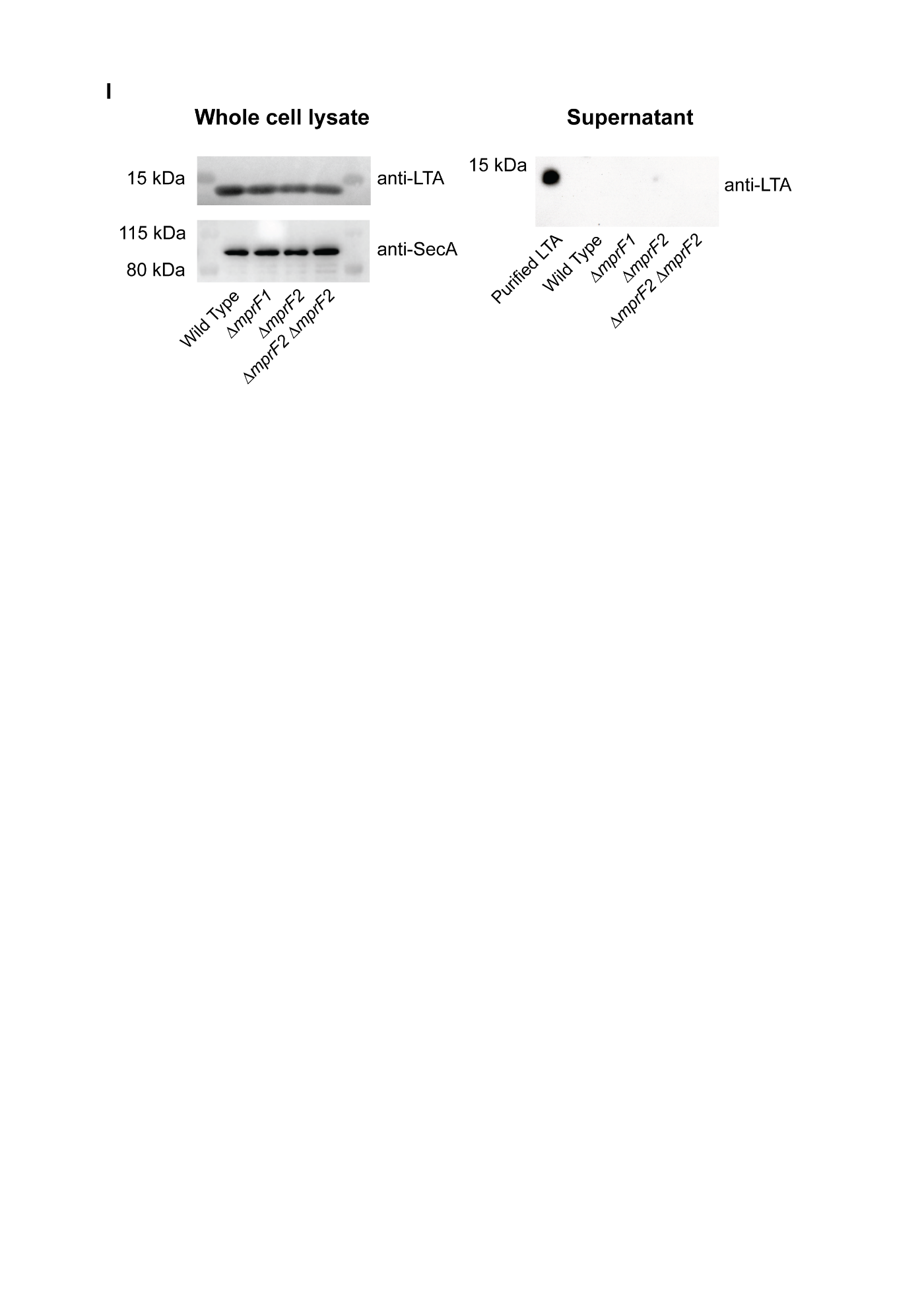

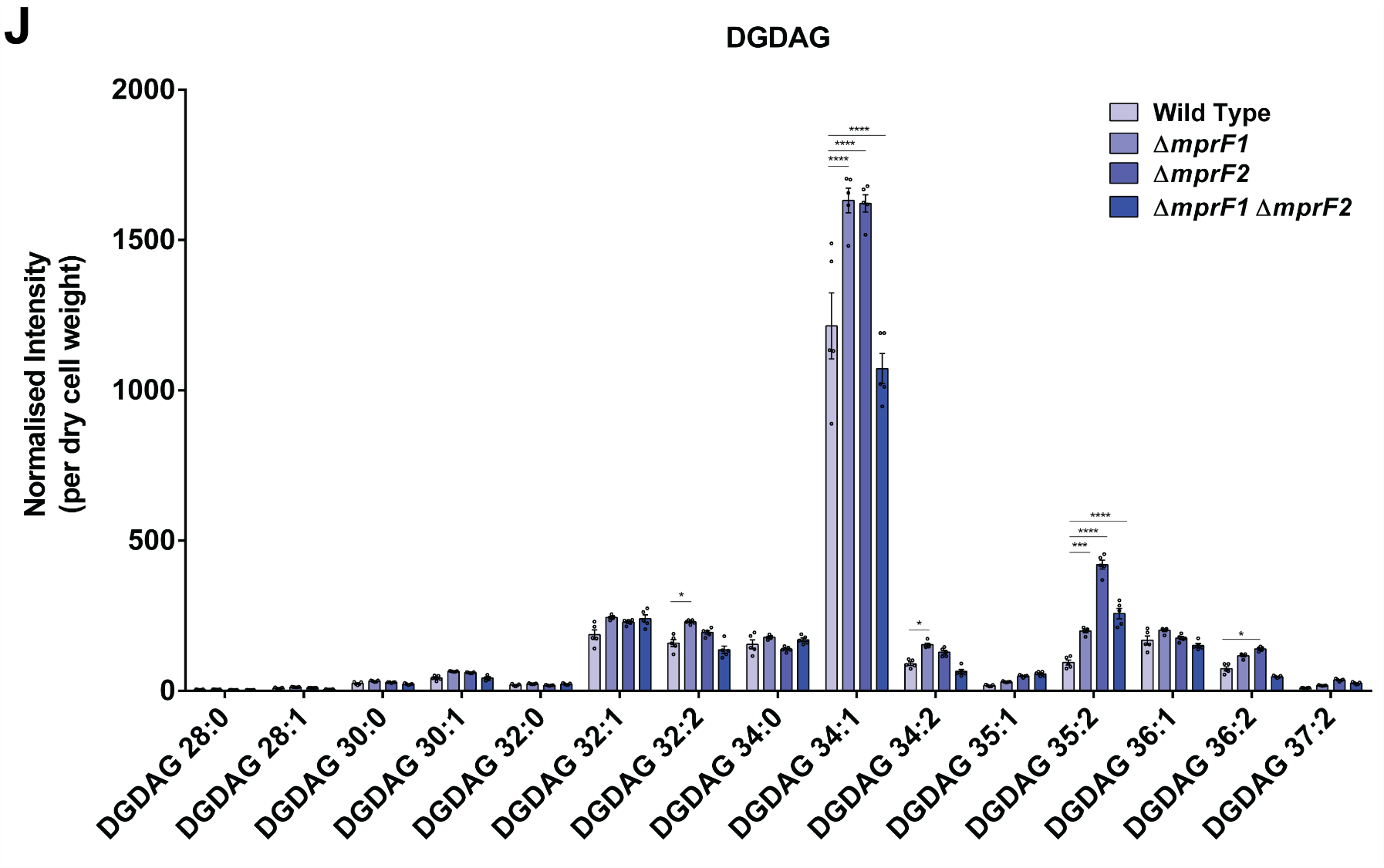

**Supplementary Fig. S1. CAMP susceptibility assessment and analysis of individual species of lysyl-PG (L-PG), phosphatidylglycerol (PG) and, diglucosyl-diacylglycerol (DGDAG) and lipoteichoic acid (LTA) in the *mprF* mutants. Deletion of *mprF* paralogs increases susceptibility to CAMPs to different degrees.** CFU enumerated after 2-hour exposure to increasing concentrations of **(A)** hBD2 or **(B)** LL-37. Error bars represent the standard error of mean from 2 biological replicates averaged from 6 technical replicates each. *, 0.05>p≥ 0.01; ****, p<0.0001; Fisher’s least significant difference (LSD) test for ANOVA. **Loss of *mprF* contributes to changes in individual species of L-PG and PG.** Normalized quantities of **(C)** 8 native lysyl-PG species, **(D)** 18 native PG species. Error bars represent the standard error of mean from 5 biological replicates. *, p<0.05; **, p<0.01; ***, p<0.001; ****, p<0.0001; Dunnett’s test for ANOVA. Inactive MprF2 does not complement L-PG and PG levels**.** Normalized amounts of total lysyl-phosphatidylglycerol (L-PG) **(E)** and total phosphatidylglycerol (PG) quantities **(F)** in *E. faecalis* wild type and *mprF* mutants are shown. Each bar represents the mean ± standard error of measurement calculated from 5 biological replicates, each represented by an open circle. **, p<0.001; ****, p<0.0001; Fisher’s least significant difference (LSD) test for ANOVA. **Inactive MprF2 (possessing the D731A, R734S catalytic domain inactivating mutations) does not complement L-PG and PG levels at the individual species level.** Normalized quantities of **(G)** 8 native lysyl-PG (L-PG) species, and **(H)** 18 native phosphatidylglycerol (PG) species. Each bar represents the mean ± standard error of measurement calculated from 5 biological replicates, each represented by an open circle. **No difference in lipoteichoic acid (LTA) levels between wild type and *mprF* mutants. (I)** Western blots for LTA in whole cell lysates and supernatants. SecA used as a loading control. **Loss of *mprF* contributes to changes in individual species of DGDAG.** Semi-quantitative analysis of **(J)** 15 native DGDAG species in the *mprF* mutants. Error bars represent the standard error of the mean from 5 biological replicates. *, p<0.05; **, p<0.01; ***, p<0.001; ****, p<0.0001; Dunnett’s test for ANOVA.

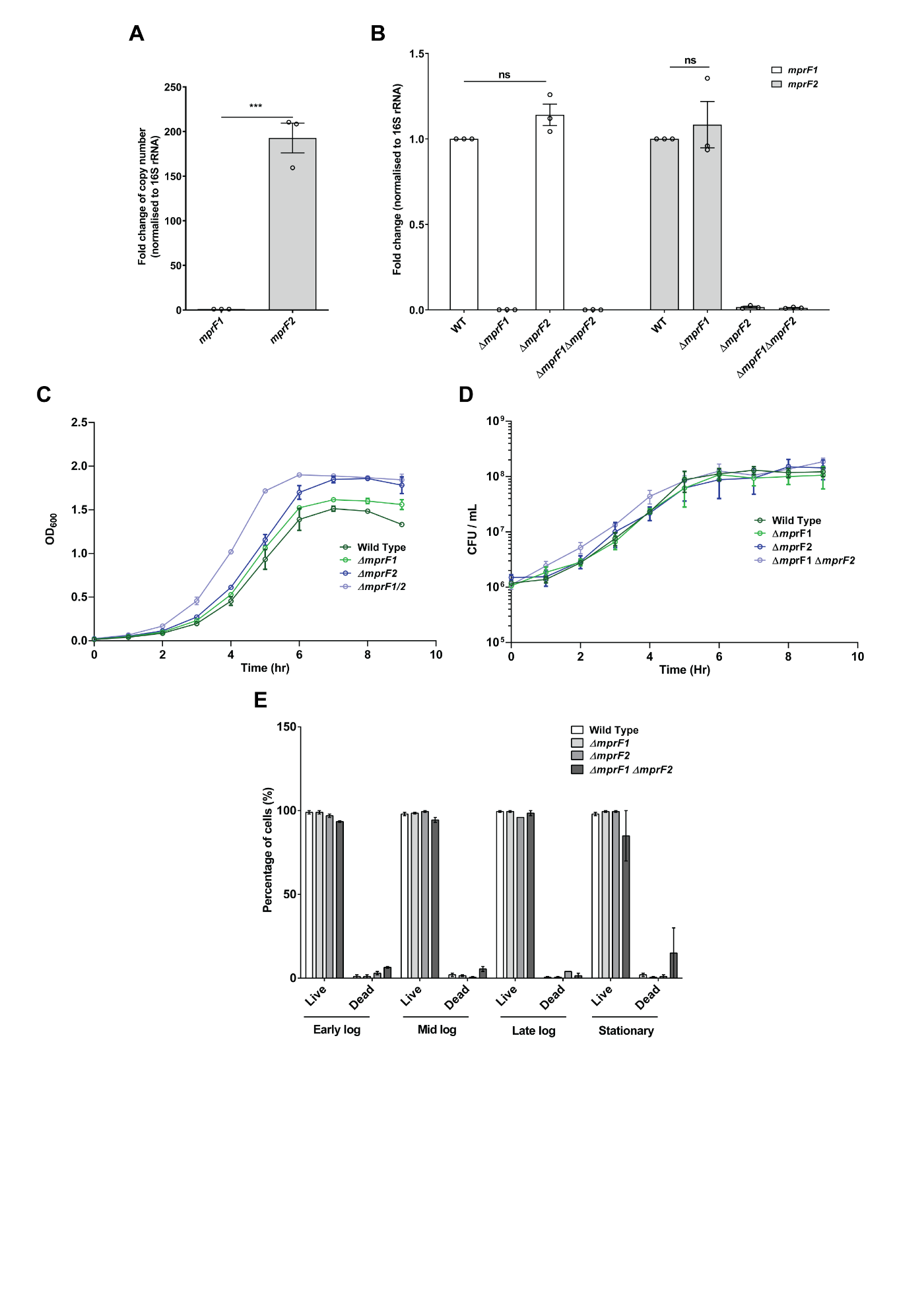

**Supplementary Fig. S2. Expression of the *mprF* paralogs and viability of the *mprF* mutants. qPCR of the *mprF* paralogs reveal higher *mprF2* expression as compared to *mprF1* in the wild type, and that deletion of one *mprF* paralog does not significantly impact expression of the remaining paralog. (A)** Absolute quantitation of *mprF1* and *mprF2* transcripts using reverse transcription quantitative PCR (RT-qPCR) of mid-log phase cultures of the wild type displaying fold change of *mprF2* relative to *mprF1* copy number. **(B)** Relative quantification of *mprF1* and *mprF2* transcripts using RT-qPCR of mid-log phase cultures of WT, Δ*mprF1*, Δ*mprF2* and Δ*mprF1* Δ*mprF2.* Data displays fold change of transcripts relative to the wild type. Each bar represents the mean ± standard error of measurement calculated from 3 biological replicates averaged from 3 technical replicates each. ***, p<0.001; Unpaired t-test for (A), Fisher’s least significant difference (LSD) test for ANOVA for (B). **No difference in growth rates or cell viability between wild type and *mprF* mutants. (C)** Optical density measured at 600 nm for the wild type and *mprF* mutants across 9 time points is shown. Error bars represent the standard error of the mean from 2 biological replicates averaged from 2 technical replicates each. **(D)** CFU enumeration results of the wild type and *mprF* mutants across 9 time points are shown. Error bars represent the standard error of the mean from 2 biological replicates averaged from 5 technical replicates each. **(E)** Percentage of live and dead cells for OG1RF WT and *mprF* mutants across 4 bacterial growth phases. Enumerated from at least 100 cells per strain.

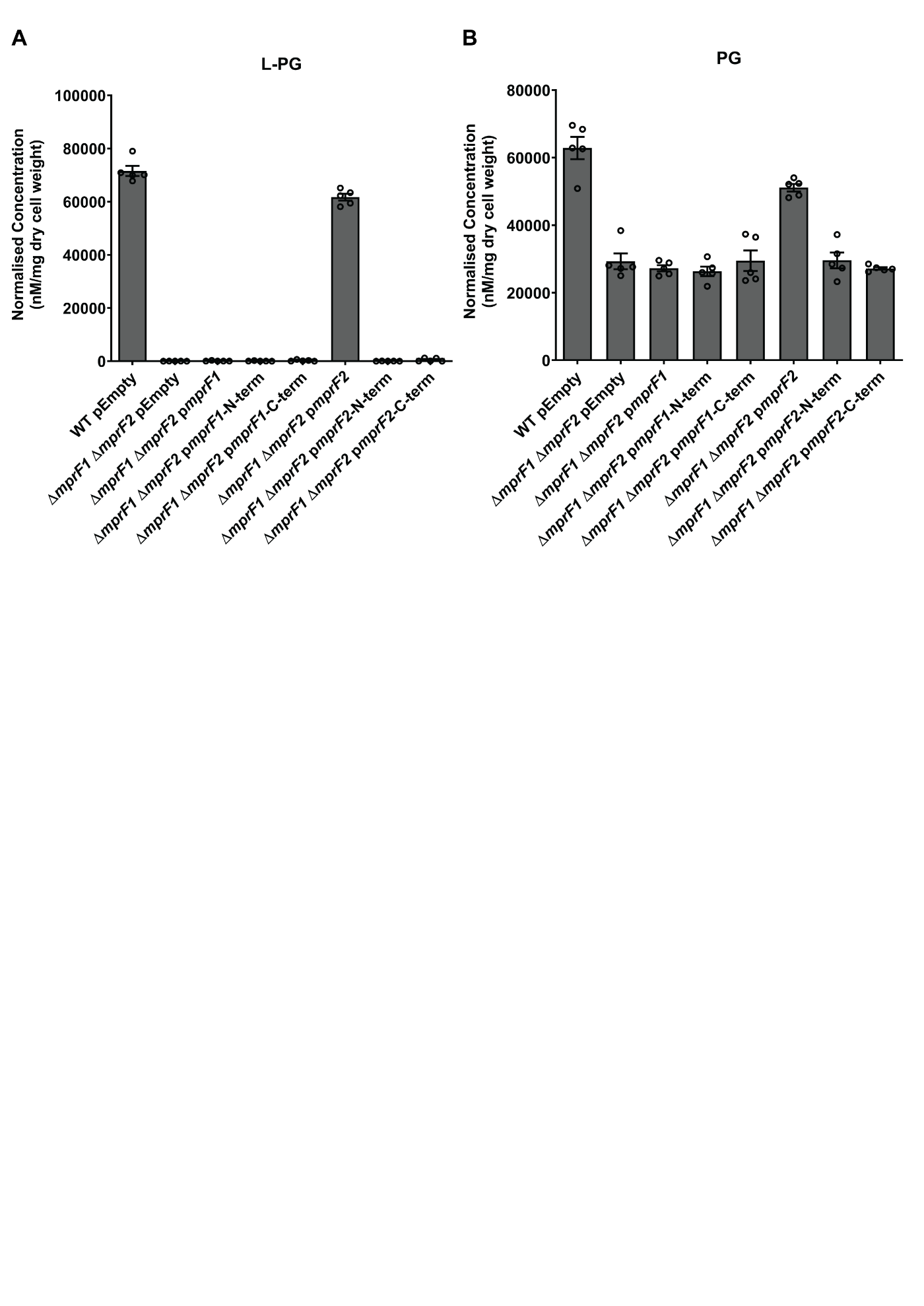

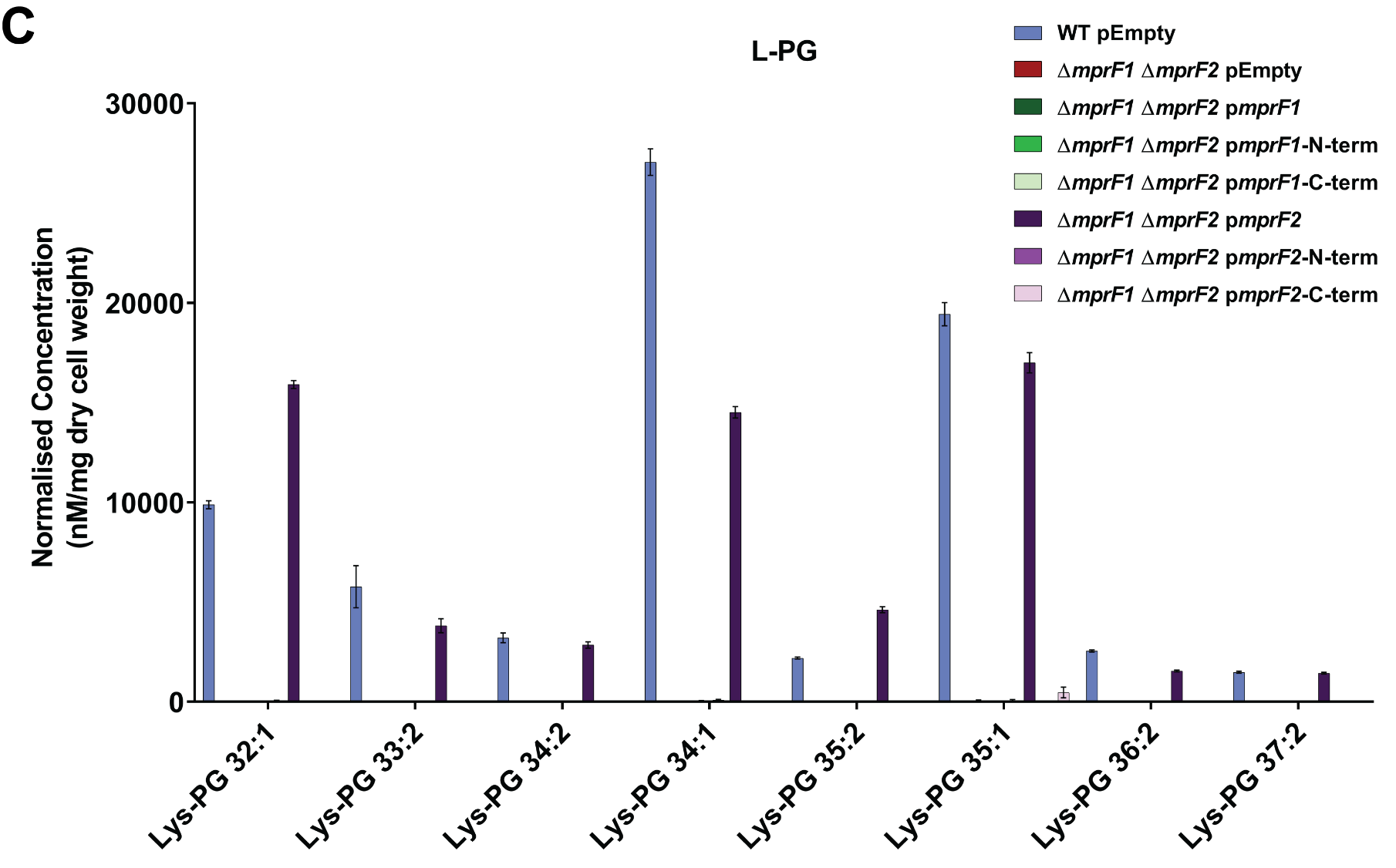

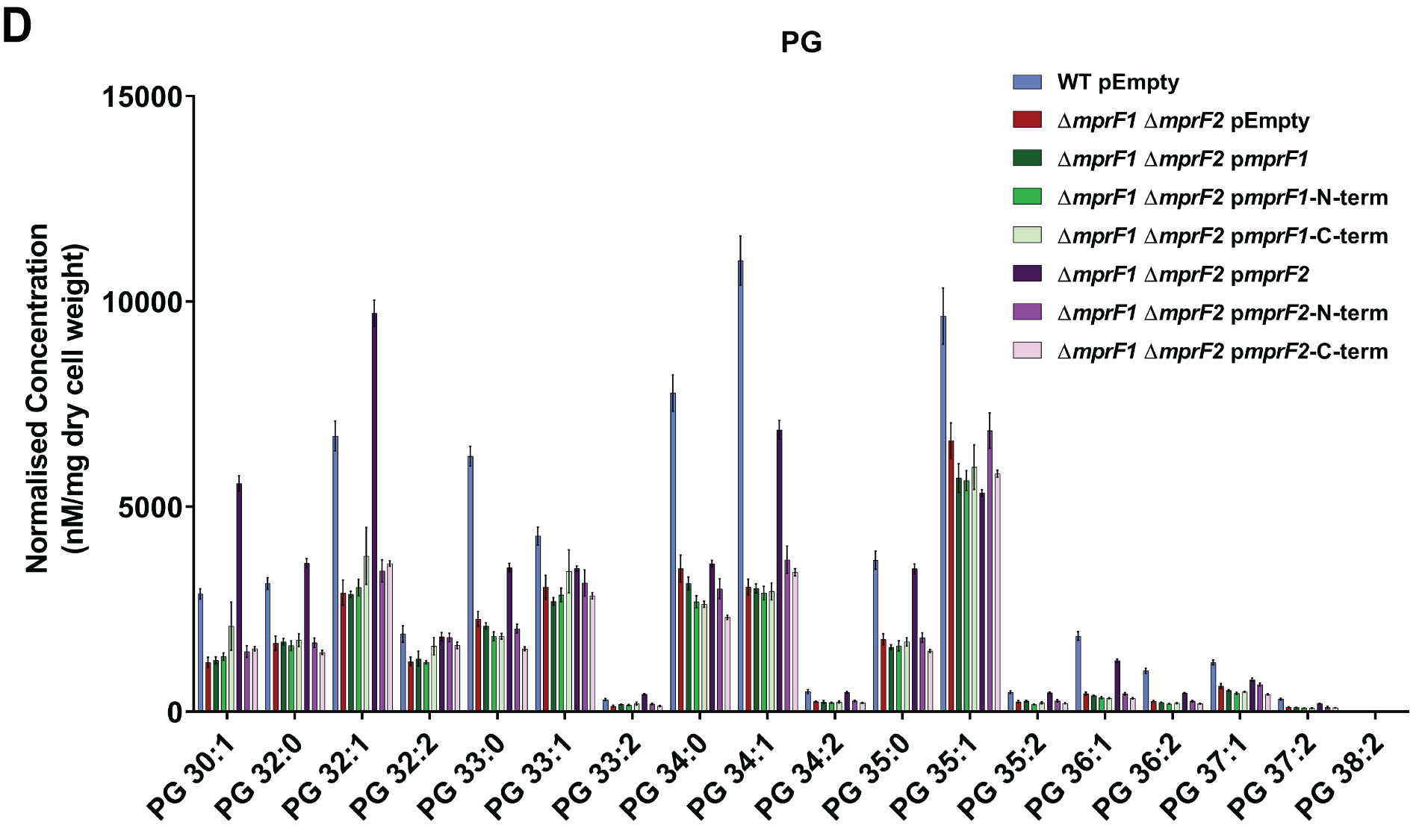

**Supplementary Fig. S3. Expression of individual *mprF* domains (N-terminal flippase and C-terminal catalytic domains)** **reveals that full length MprF is required for restoration of PG and L-PG levels in Δ*mprF1*Δ*mprF2*.** Normalized quantities of PG and L-PG from the Δ*mprF1*Δ*mprF2* with plasmid-based expression using the PsrtA promoter of full length *mprF1*, *mprF2* or the N-terminal, C-terminal domains of either *mprF*. Total normalized quantities of **(A)** L-PG and **(B)** PG in the strains tested as well as individual normalized quantities of **(C)** L-PG and **(D)** PG. Error bars represent the standard error of mean from 5 biological replicate. Plasmid backbone used here for all strains are pGCP123-P*srtA*, except for pEmpty which are pGCP123 instead.

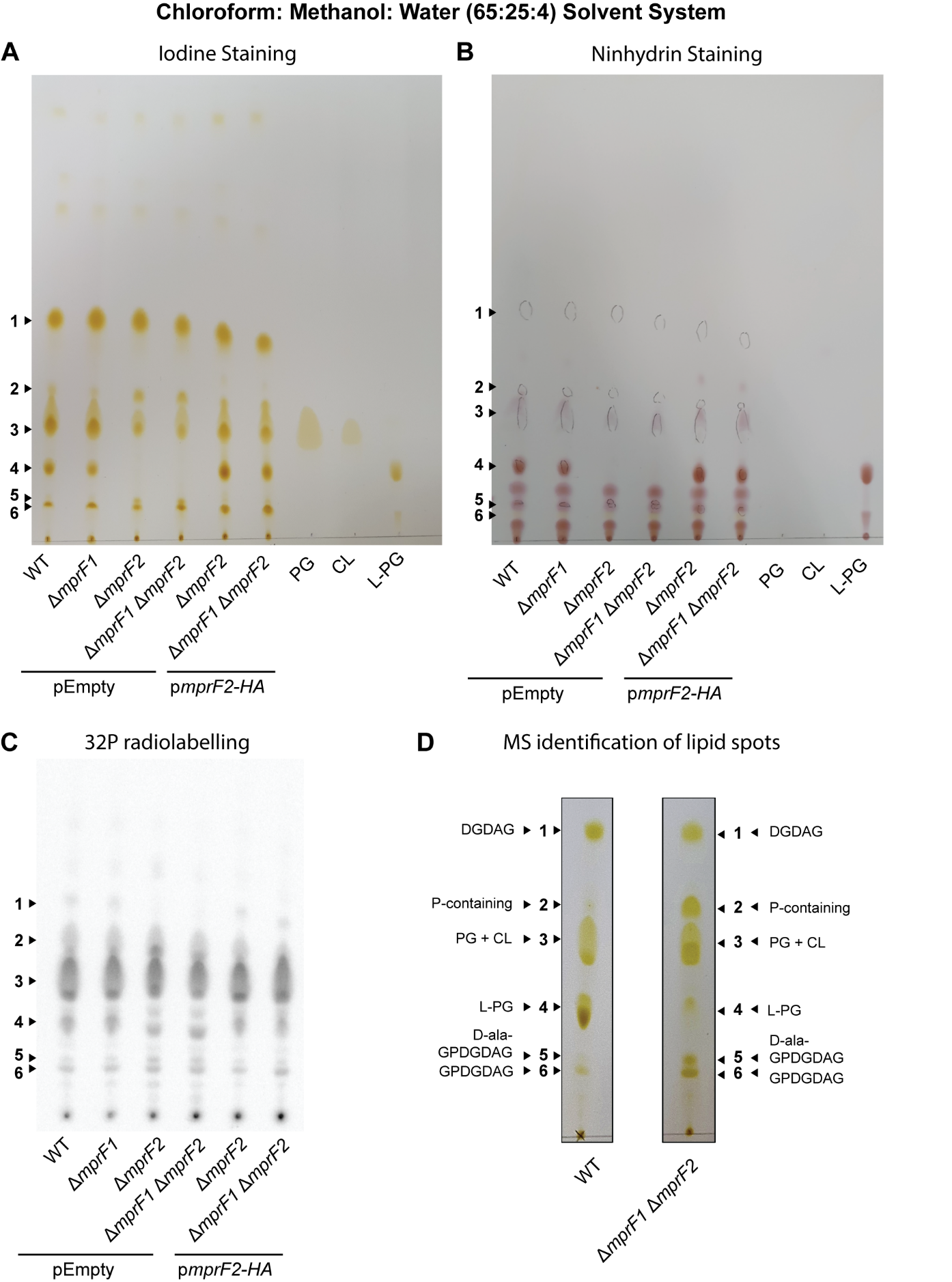

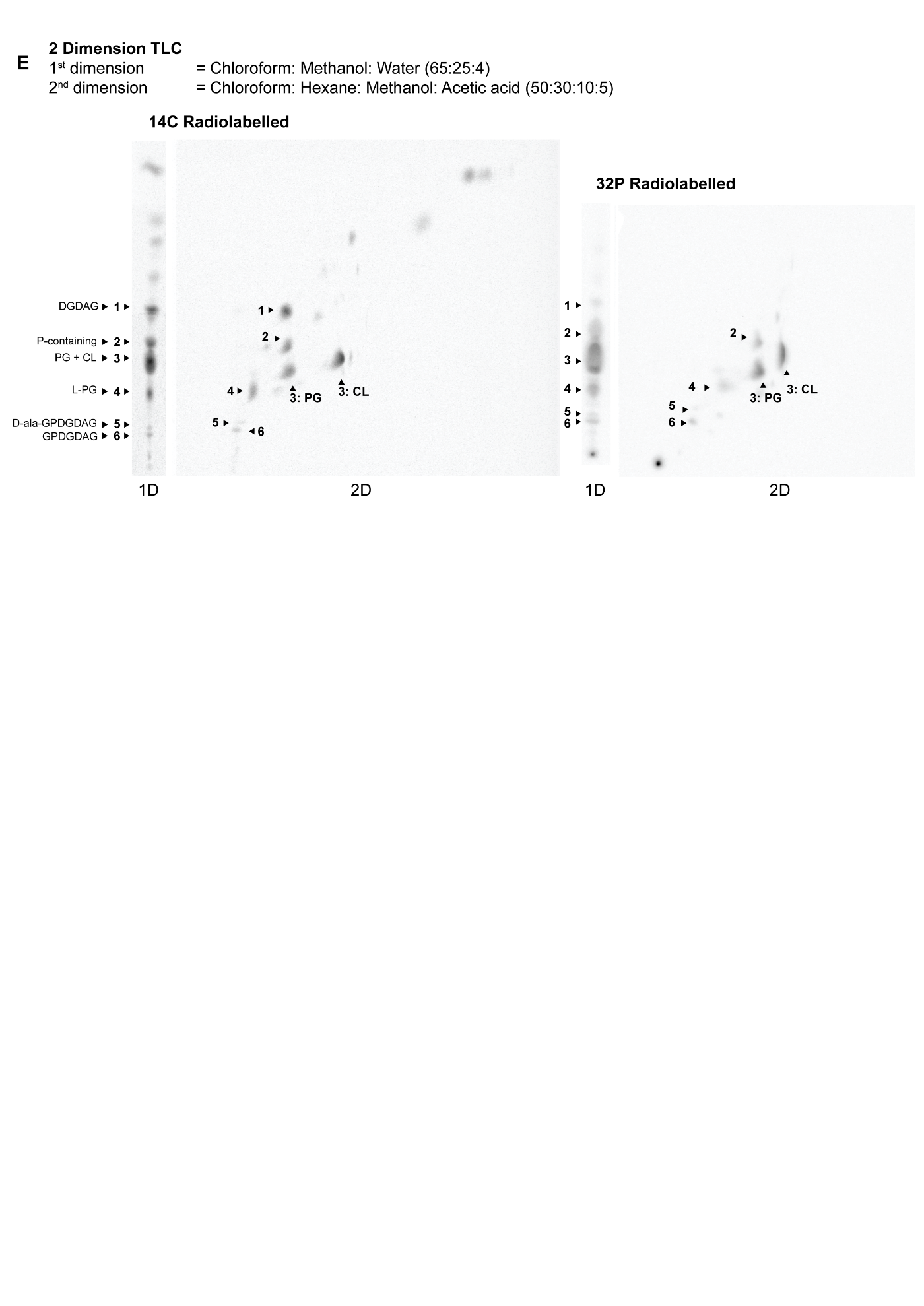

**Supplementary Fig. S4. Identification of lipid spots in 1D- and 2D-TLCs** **of lipid extracts from WT and the *mprF* mutants.** **(A)** Iodine staining, **(B)** ninhydrin staining and **(C)** 32P-radiolabeling staining of 1D-TLCs of empty vector controls of WT, ∆*mprF1*, ∆*mprF2*, ∆*mprF1*∆*mprF2* and p*mprF2*-HA complemented ∆*mprF2*, ∆*mprF1*∆*mprF2* together with PG, CL and lysyl-PG standards under chloroform: methanol: water (65:25:4) solvent system. Iodine stains most lipids while ninhydrin stains amino modified lipids. 6 major spots were observed. **(D)** These spots were scraped off iodine strained TLC plates of WT and ∆*mprF1*∆*mprF2* and placed through lipid extraction before analysis using LC-MS. Spots were identified and labelled accordingly. **(E)** 1D and 2-D TLCs of 14C and 32P labelled lipid extracts of wild type and ∆*mprF1*∆*mprF2*. chloroform: methanol: water (65:25:4) solvent system was used in the first dimension while chloroform: hexane, methanol, acetic acid (50:30:10:5) was used in the second dimension. Comparing positions of the spots in the first and second dimension, we can resolve the positions of PG and CL. In the solvent system used in the second dimension, CL migrates ahead of PG.

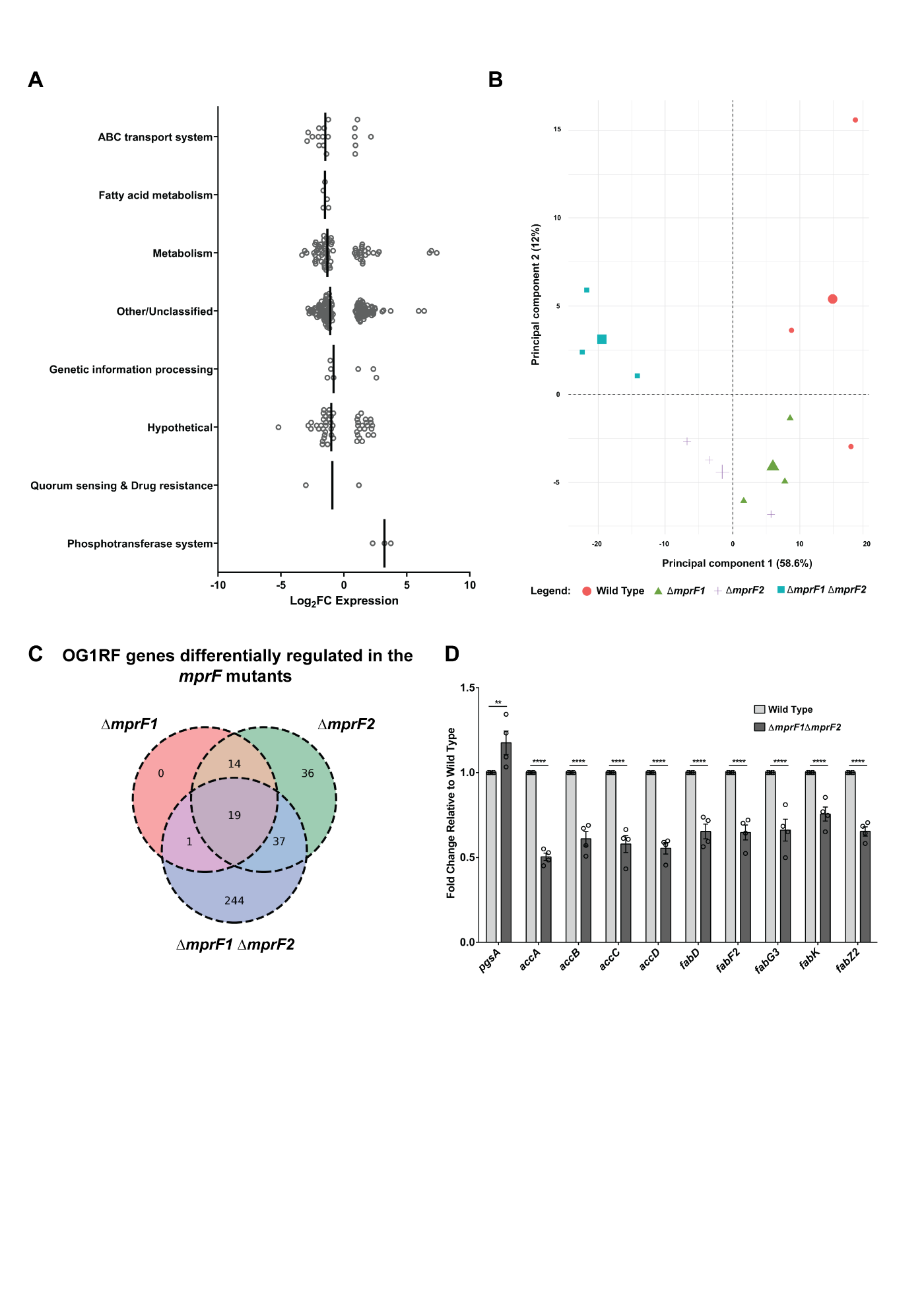

**Supplementary Fig. S5.** **Δ*mprF2* is more similar to Δ*mprF1*Δ*mprF2* in terms of transcriptional gene expression than Δ*mprF1*. (A)** Global gene expression profile comparing *E. faecalis* OG1RF with Δ*mprF1*Δ*mprF2* (Log_2_FC expression; FDR<0.05)*.* Gene functional categories are based on KEGG pathway membership and manually curated for category assignment. The majority of the differentially expressed genes were categorized under metabolism and membrane transport. A vertical black line represents the median value for each category. **(B)** Principal component analysis of global gene expression profile comparing *E. faecalis* wild type (orange circle), Δ*mprF1* (green triangle), Δ*mprF2* (blue square), and Δ*mprF1*Δ*mprF2* (purple cross). Smaller symbols represent the spread of each biological replicate and larger symbols represent the centroid of their respective clusters. The two dimensions measured account for 70.6% of the variability of all differentially expressed genes. **(C)** Venn diagram represents differentially expressed *E. faecalis* genes in the *mprF* mutants relative to wild type. The number of genes (FDR<0.05) in each section was calculated from Excel Table S1D-F, where intersections of genes were determined by matching locus tags. **(D)** RT-qPCR of mid-log phase cultures of wild type and ∆*mprF1*∆*mprF2* showing a slight increase in *pgsA* and decrease in fatty acid biosynthesis gene expression levels. Each bar represents the mean ± standard error of measurement calculated from 4 biological replicates averaged from 3 technical replicates each. **, p<0.01; ****, p<0.0001; Fisher’s least significant difference (LSD) test for ANOVA.

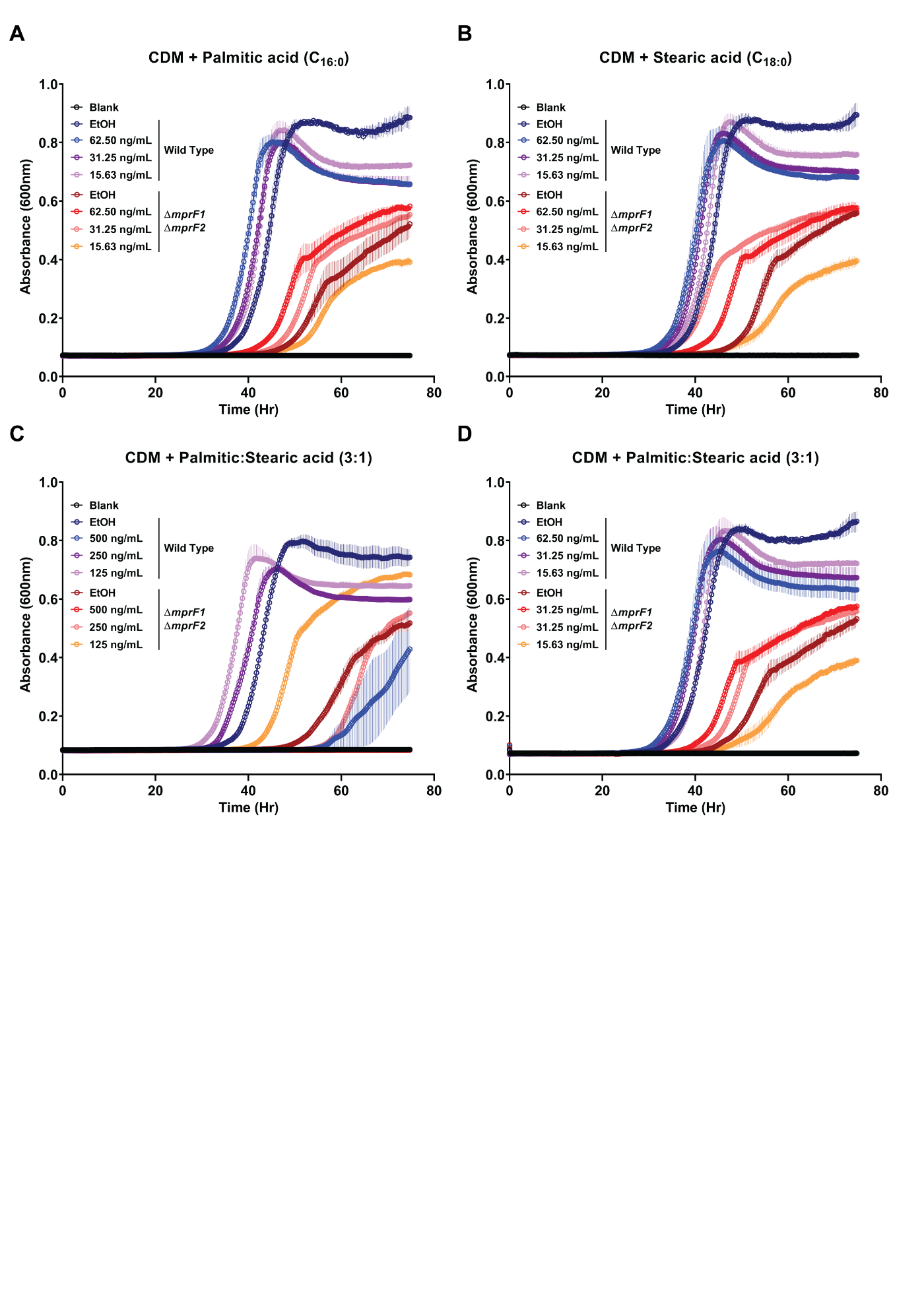

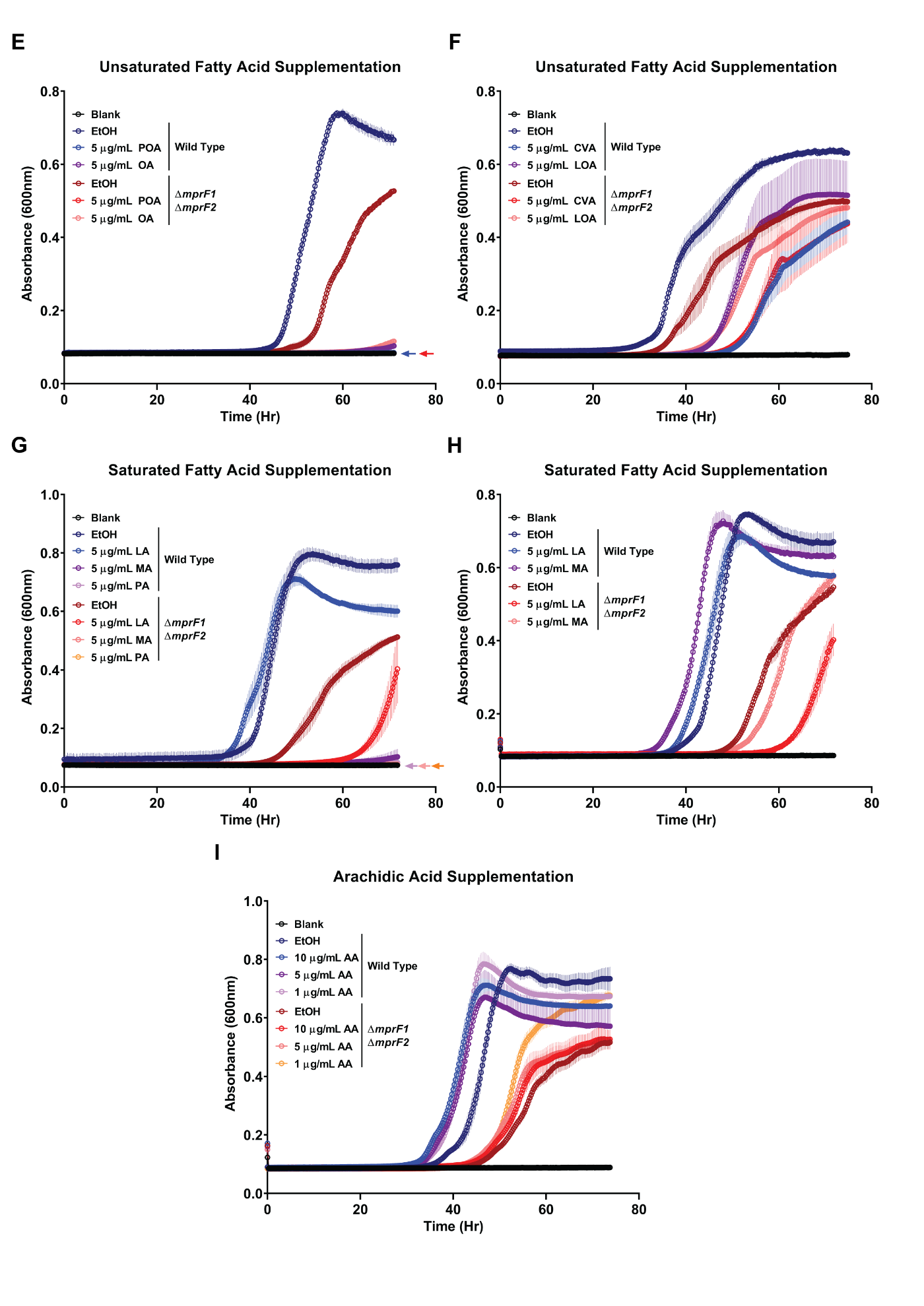

**Supplementary Fig. S6. Saturated fatty acid supplementation at low concentrations promotes growth of *mprF* mutants cultured in chemically defined media (CDM) while high concentrations of unsaturated fatty acids, myristic acid, and lauric acid fail to promote growth.** Growth curves of wild type and ∆*mprF1*∆*mprF2* grown under fatty acid supplementation of **(A)** palmitic acid (C_16:0_) or **(B)** stearic acid (C_18:0_) at lower concentrations of 62.5, 31.25 or 15.63 ng/ml. **(C, D)** Growth curves of combinations of palmitic and stearic acid in a 3:1 ratio at varying concentrations. Growth curves of wild type and ∆*mprF1*∆*mprF2* grown under higher concentrations of unsaturated fatty acid supplementation of **(E)** palmitoleic acid (C_16:1 cis-9_) or oleic acid (C_18:1 cis-9_) and **(F)** cis-vaccenic acid (C_18:1 cis-7_) or linoleic acid (C_18:2 cis-9,12_), as well as **(G)** lauric acid (C_12:0_), myristic acid (C_14:0_) or palmitic acid (C_16:0_) and **(H)** stearic acid (C_18:0_) or arachidic acid (C_18:0_). Fatty acids were supplemented at 5 μg/ml. **(I)** Growth curves of 1, 5, and 10 μg/ml of arachidic acid. Equal volumes of ethanol (EtOH) were used as the solvent control for comparison. Total fatty acid concentrations are shown. Data points for all graphs represents the mean ± standard error of measurement calculated from 3 biological replicates averaged from 3 technical replicates each. Fatty acid abbreviations: [unsaturated fatty acids – POA: palmitoleic acid (C_16:1 cis-9_), OA: oleic acid (C_18:1 cis-9_) CVA: cis-vaccenic acid (C_18:1 cis-7_), LOA: linoleic acid (C_18:2 cis-9,12_)] and [saturated fatty acids – LA: lauric acid (C_12:0_), MA: myristic acid (C_14:0_), PA: palmitic acid (C_16:0_), SA: stearic acid (C_18:0_), AA: arachidic acid (C_18:0_)]. For curves that overlap the blank, an arrow corresponding to their respective colors are shown next to the blank curve.

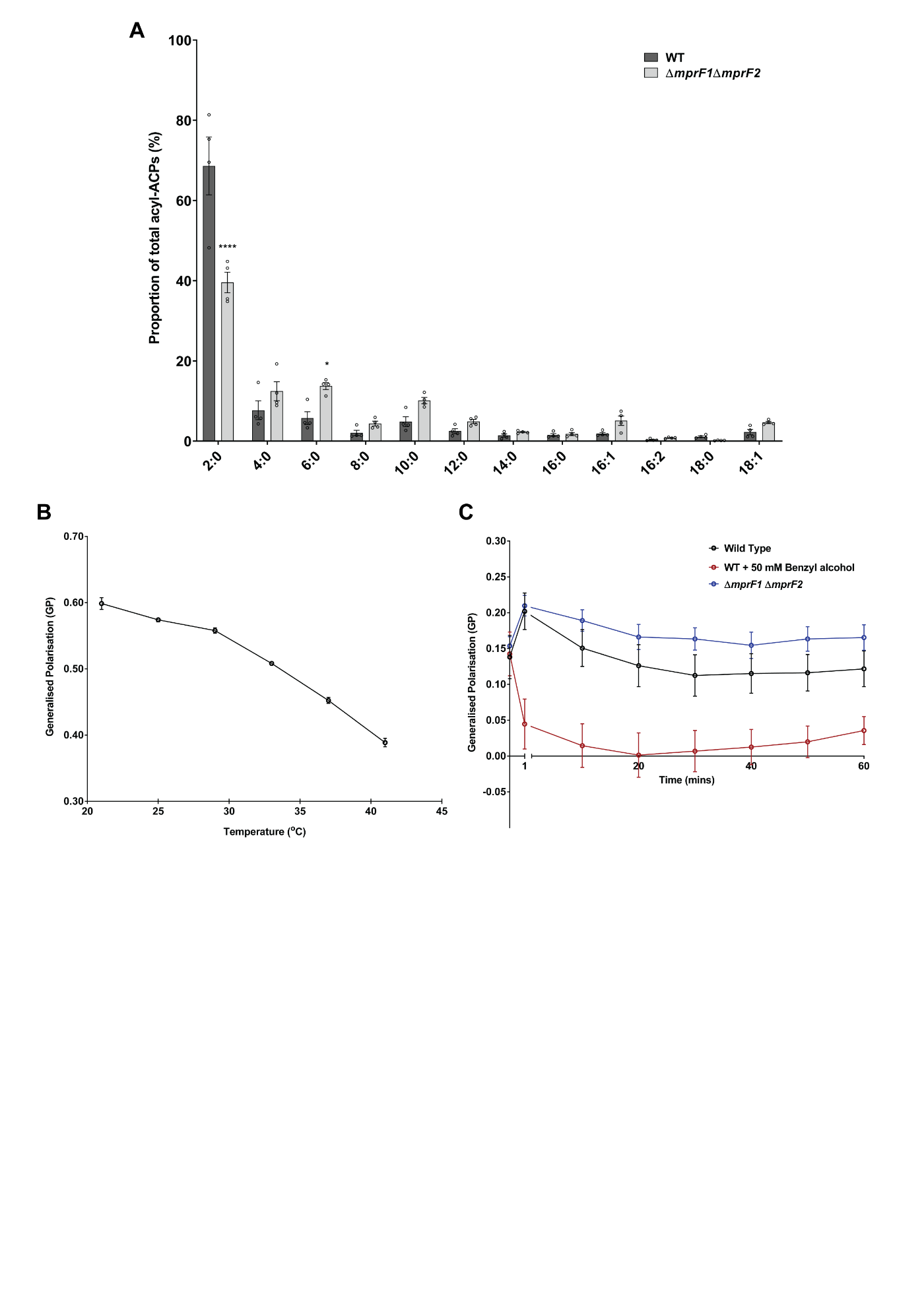

**Supplementary Fig. S7. Acyl-ACP analysis and control experiments testing sensitivity of the Laurdan dye. Analysis of individual species of acyl-ACPs in WT and Δ*mprF1*Δ*mprF2*. (A)** Proportion of acyl-ACPs within WT and Δ*mprF1*Δ*mprF1*. Values obtained through mass spectrometry were normalized against internal standards and total protein concentration before conversion to proportion. Error bars represent the standard error of mean from 4 biological replicates. *, p<0.05; **, p<0.01; ****, p<0.0001; Fisher’s LSD test for ANOVA. **Control experiments show that the Laurdan assay can sensitively and accurately measure differences in membrane fluidity.** Wild type cells grown at a range of temperatures that result in different membrane fluidities **(B)** and exposure to the membrane fluidizer, benzyl alcohol **(C)**. Benzyl alcohol was spiked in just after the initial reading at time, t = 0 minutes. Experiments are done in a microtiter plate set-up with measurements taken using a plate reader instead due to the technical constraints in the microscope set up in **Fig. 7C**. Each bar represents the mean ± standard error of measurement calculated from 3 biological replicates.
