## Supplementary Tables S1A-S1G for "Depleting cationic lipids involved in antimicrobial resistance drives adaptive lipid remodeling in *Enterococcus faecalis*"

**Table S1 - Supplementary tables for methods**

**Table S1A.** Bacteria Strains and Culture Conditions

| **Bacterial Strains** | **Relevant Information / Genotype** | **References / Source** | **Media Used** |
| --- | --- | --- | --- |
| ***E. coli*** | | | |
| Stellar^TM^ Competent Cells (*E. coli* HST08) | *F–, endA1, supE44, thi-1, recA1, relA1, gyrA96, phoA, Φ80d lacZΔM15, Δ(lacZYA - argF) U169, Δ(mrr - hsdRMS - mcrBC), ΔmcrA, λ–* | Clontech, Takara Bio Inc., Japan | - |
| DH5α pCYW2 | DH5α harbouring pCYW2*; Kan^r^  *pCYW2: pGCP123 parent plasmid (5) with PsrtA promoter. Expression vector. | Lab stock | LB, Miller broth (BD, USA) with 50 µg/mL kanamycin (Thermo Scientific, USA) |
| DH5α pMSP3535 | DH5α harbouring pMSP3535*; Erm^r^  *pMSP3535: Shuttle vector for nisin-controlled inducible expression. | pMSP3535 was a gift from Gary Dunny, University of Minnesota (Addgene plasmid # 46886)  (6) | LB, Miller broth (BD, USA) with 300 µg/mL erythromycin (Sigma Alrdrich, USA) |
| Stellar^TM^ Competent Cells pMSP3545-d*cas9* | DH5α harbouring pMSP3545-d*cas9**; Erm^r^  *pMSP3535-dcas9: Shuttle vector for nisin-inducible expression of dcas9 | Constructed by Irina Afonina (7) | LB, Miller broth (BD, USA) with 300 µg/mL erythromycin (Sigma Alrdrich, USA) |
| DH5α pGCP123-*mprF1-HA* | DH5α harbouring pGCP123-*mprF1*-HA*  * pGCP123-*mprF1*-HA: Shuttle expression vector for *mprF1* complementation | This study | LB, Miller broth (BD, USA) with 50 µg/mL kanamycin (Thermo Scientific, USA) |
| DH5α pGCP123-*mprF2-HA* | DH5α harbouring pGCP123-*mprF2*-HA*  * pGCP123-*mprF2*-HA*:* Shuttle expression vector for mprF2 complementation |  |  |
| DH5α pGCP123-*mprF2* | DH5α harbouring pGCP123-*mprF2**  * pGCP123-*mprF2:* Shuttle expression vector for mprF2 complementation |  |  |
| DH5α pGCP123-P*srtA* | DH5α harbouring pGCP123-P*srtA**  *pGCP123-*mprF2*-HA*:* Shuttle expression vector for P*srtA* directed expression | This study | LB, Miller broth (BD, USA) with 50 µg/mL kanamycin (Thermo Scientific, USA) |
| ***E. faecalis*** | | | |
| OG1X | Strep^r^, wild type | Lab stock | BHI (Neogen, USA) |
| OG1RF | Fus^r^, Rif^r^, wild type strain | American Type Culture Collection (ATCC^©^ 47077™) |  |
| OG1RF *∆mprF1* | Fus^r^, Rif^r^, *∆mprF1* | (2) |  |
| OG1RF *∆mprF2* | Fus^r^, Rif^r^, *∆mprF2* | (2) |  |
| OG1RF *∆mprF1∆mprF2* | Fus^r^, Rif^r^, *∆mprF1∆mprF2* | This study |  |
| OG1RF *∆mprF1 pGCP123-mprF1-HA* | Fus^r^, Rif^r^, *∆mprF1 pGCP123-mprF1-HA* | This study | BHI (Neogen, USA) with 500 µg/mL kanamycin (Thermo Scientific, USA) |
| OG1RF *∆mprF2 pGCP123-mprF2-HA* | Fus^r^, Rif^r^, *∆mprF2 pGCP123-mprF2-HA* | This study |  |
| OG1RF *∆mprF1∆mprF2 pGCP123-mprF1-HA* | Fus^r^, Rif^r^, *∆mprF1∆mprF2 pGCP123-mprF1-HA* | This study |  |
| OG1RF *∆mprF1∆mprF2 pGCP123-mprF2-HA* | Fus^r^, Rif^r^, *∆mprF1∆mprF2 pGCP123-mprF2-HA* | This study |  |
| OG1RF *∆mprF2 pGCP123-mprF2* | Fus^r^, Rif^r^, *∆mprF2 pGCP123-mprF2* | This study |  |
| OG1RF *∆mprF1∆mprF2 pGCP123-mprF2* | Fus^r^, Rif^r^, *∆mprF1∆mprF2 pGCP123-mprF2* | This study |  |
| OG1RF *∆mprF1∆mprF2 pGCP123-mprF2 (D731A, R734S)**  *also written as p*mprF2*-inactive | Fus^r^, Rif^r^, *∆mprF2 pGCP123-mprF2 (D731A,R734S)* | This study |  |
| OG1RF *∆mprF1∆mprF2 pGCP123-mprF2 (D731A, R734S)**  *also written as p*mprF2*-inactive | Fus^r^, Rif^r^, *∆mprF1∆mprF2 pGCP123-mprF2 (D731A,R734S)* | This study |  |
| OG1RF *pABG5* | Fus^r^, Rif^r^, *pABG5* | This study | BHI (Neogen, USA) with 500 µg/mL kanamycin (Thermo Scientific, USA) |
| OG1RF *∆mprF1 pABG5* | Fus^r^, Rif^r^, *∆mprF1*, *pABG5* |  |  |
| OG1RF *∆mprF2 pABG5* | Fus^r^, Rif^r^, *∆mprF2*, *pABG5* |  |  |
| OG1RF *∆mprF1∆mprF2 pABG5* | Fus^r^, Rif^r^, *∆mprF1∆mprF2* *pABG5* |  |  |
| OG1RF  *∆mprF1∆mprF2*  p*mprF1* | Fus^r^, Rif^r^, *∆mprF1∆mprF2* pGCP123-P*srtA*-*mprF1* | This study | BHI (Neogen, USA) with 500 µg/mL kanamycin (Thermo Scientific, USA) |
| OG1RF  *∆mprF1∆mprF2*  p*mprF1*-N-term | Fus^r^, Rif^r^, *∆mprF1∆mprF2* pGCP123-P*srtA*-*mprF1-*N-term |  |  |
| OG1RF  *∆mprF1∆mprF2*  p*mprF1­*-C-term | Fus^r^, Rif^r^, *∆mprF1∆mprF2* pGCP123-P*srtA*-*mprF1*-C-term |  |  |
| OG1RF  *∆mprF1∆mprF2*  p*mprF2* | Fus^r^, Rif^r^, *∆mprF1∆mprF2* pGCP123-P*srtA*-*mprF2* |  |  |
| OG1RF  *∆mprF1∆mprF2*  p*mprF2*-N-term | Fus^r^, Rif^r^, *∆mprF1∆mprF2* pGCP123-P*srtA*-*mprF2-*N-term |  |  |
| OG1RF  *∆mprF1∆mprF2*  p*mprF12*-C-term | Fus^r^, Rif^r^, *∆mprF1∆mprF2* pGCP123-P*srtA*-*mprF2*-C-term |  |  |

**Table S1B.** Lipid standards used. Procured from Avanti Polar Lipids.

| **Standard** | **Usage** | **Catalogue #** | **Stock conc. [mg/ml]** | **Final conc. [µg/ml]** | **Precursor ion (m1) m/z** | **Fragment ion (m3) m/z** |
| --- | --- | --- | --- | --- | --- | --- |
| PG 14:0 | Internal | 840445P | 1 | 5 | 665.5 [M-H]^-^ | 153 |
| L-PG 16:0 | Internal | 840520P | 0.1 | 4 | 849.6 [M-H]^-^ | 145 |
| MGDAG 34:1 | External | 840522P | 1 | variable | 774.6 [M+NH_4_]^+^ | 313 |

**Table S1C.** Primers used for RT-qPCR

| **Primer** | **Target** | **Gene Type** | **Sequence (5’ 🡪 3’)** | **Tm (^o^C)** |
| --- | --- | --- | --- | --- |
| gyrA_F | *gyrA* | Housekeeping | TGTTCGTCGGGATGTGAGTG | 55 |
| gyrA_R | *gyrA* | Housekeeping | GGTACGCCTTTTTCGATGGC | 55 |
| 16S_F | 16s rRNA | Housekeeping | GGATAACACTTGGAAACAGG | 55 |
| 16S_R | 16s rRNA | Housekeeping | TCCTTGTTCTTCTCTAACAA | 55 |
| pgsA_F | *pgsA* | Target | CGACACAACGTTAGCCGTTA | 55 |
| pgsA_R | *pgsA* | Target | AAGGCGGTCATCACTAGCAT | 55 |
| accA_F | *accA* | Target | GCGATACCCTTCAGGATTAG | 55 |
| accA_R | *accA* | Target | CTTTGCCGATGATTTAGCTG | 55 |
| accB_F | *accB* | Target | GCTTCAACGATACACACAACG | 55 |
| accB_R | *accB* | Target | ACCGACAACCAATGAAAAGA | 55 |
| accC_F | *accC* | Target | CGATATGACGTGCTGGATAA | 55 |
| accC_R | *accC* | Target | TACCCAGTGATGTTAAAGGCA | 55 |
| accD_F | *accD* | Target | GTTGGATCAGTCAATACCGTAAG | 55 |
| accD_R | *accD* | Target | TGATTTTCACTGCATCTGGTG | 55 |
| fabD_F | *fabD* | Target | TCACGATTTGTTGTGGTGTAT | 55 |
| fabD_R | *fabD* | Target | GTCAGTACATGACAGAAGCAGC | 55 |
| fabF2_F | *fabF2* | Target | CTGGTGTTGGTGATGTCATA | 55 |
| fabF2_R | *fabF2* | Target | TGGATTTGTGATGGGAGAAGG | 55 |
| fabG3_F | *fabG3* | Target | TATCATTCGTAATCCCAGCG | 55 |
| fabG3_R | *fabG3* | Target | ATTGAAGCCTTTGGGGTAAAAT | 55 |
| fabK_F | *fabK* | Target | TCAATGATTTCTTGGGCTGT | 55 |
| fabK_R | *fabK* | Target | TCCAAAAAGAAGTGCCTGAT | 55 |
| fabZ2_F | *fabZ2* | Target | CCCTTTGAATTCAGGCATTG | 55 |
| fabZ2_F | *fabZ2* | Target | GCGTTGTAGCGAAAAAGAATG | 55 |
| mprF1_F | *mprF1* | Target | CCAGCGCTCTGGTTTTTCAC | 55 |
| mprF1_R | *mprF1* | Target | AGCAAACATCCCACCTAGCC | 55 |
| mprF2_F | *mprF2* | Target | ATGATGGTGCTCCTAGCCAC | 55 |
| mprF2_R | *mprF2* | Target | CAGCAATCCCACGTCCCATA | 55 |

**Table S1D.** Components in chemically defined media (CDM)

| **Component** | **g/L** | **Component** | **g/L** |
| --- | --- | --- | --- |
| KH_2_PO_4_ | 3.6000 | D-glucose | 15.0000 |
| NaCl | 3.0000 | L-histidine | 0.1700 |
| (NH_4_)_2_SO_4_ | 1.0000 | L-isoleucine | 0.2400 |
| MOPS | 13.0500 | L-leucine | 1.0000 |
| K-acetate | 0.9000 | L-methionine | 0.0600 |
| (NH_4_)_6_Mo_7_O_24_ **·** 4H_2_O | 0.0002 | L-valine | 0.7000 |
| ZnSO_4_ **·** 7H_2_O | 0.0050 | L-arginine | 0.7200 |
| CoCl_2_ **·** 6H_2_O | 0.0002 | L-glutamic acid | 0.7200 |
| CuSO_4_ **·** 5H_2_O | 0.0002 | Glycine | 0.3600 |
| H_3_BO_3_ | 0.0008 | L-serine | 0.6000 |
| K_2_SO_4_ | 0.0230 | L-threonine | 0.6000 |
| KI | 0.0001 | MgSO_4_ **·** 7H_2_O | 1.0000 |
| L-tryptophan | 0.2400 | CaCl_2_ **·** 2H_2_O | 0.0400 |
| L-cysteine HCl | 0.2400 | Inositol | 0.0020 |
| Ca-pantothenate | 0.0012 | Biotin | 0.0060 |
| Pyridoxal HCl | 0.0048 | Niacin | 0.0009 |
| Riboflavin | 0.0009 | p-aminobenzoic acid | 0.0001 |
| Folic acid | 0.0006 | Tricine | 1.3050 |
| Thiamine-HCl | 0.0006 | Glutathione | 0.0150 |
|  |  | Iron (II) sulfate heptahydrate | 0.0040 |

**Table S1E.** Antibody and developing solution concentrations for Western blot

| **Target of interest** | **Primary Antibody Dilution; Host** | **Secondary Antibody Dilution; Host** | **Developing Solution; Dilution Ratios (Luminol:Peroxide:Water)** |
| --- | --- | --- | --- |
| LTA | Mouse anti-LTA 1:1000 (Hycult Biotech, Netherlands) | Goat anti-mouse HRP 1:5000 (Thermoscientific, USA) | SuperSignal™ West Femto Maximum Sensitivity Substrate 1:1:8 |
| 2HA | Mouse anti-HA 1:1000 (Thermoscientific, USA) | Goat anti-mouse HRP 1:5000 (Thermoscientific, USA) | SuperSignal™ West Femto Maximum Sensitivity Substrate 1:1:8 |
| SecA | Rabbit anti-SecA 1:3000 (8) | Goat anti-rabbit HRP 1:6000 (Thermoscientific, USA) | SuperSignal™ West Femto Maximum Sensitivity Substrate 1:1:8 |

**Table S1F.** PCR Primers

| **Primer** | **Target(s)** | **Sequence** | **Purpose** |
| --- | --- | --- | --- |
| pmprF1-HA_F | genomic DNA | GCTTGATATCGAATTCATATCCTGCACGCACAACGA | To create insert for ligation into pGCP123 linearized vector |
| pmprF1-HA_R | genomic DNA | GTGGATCCCCCGGGCTGCAGAGATAAACAACCGCCACTCT |  |
| pmprF2-HA_F | genomic DNA | TTGATATCGAATTCCTGCAGTATCGTGGCTCTGTTATTTG | To create insert for ligation into pGCP123 linearized vector |
| pmprF2-HA_R | genomic DNA | TAATAAAAAATCGAGCATGCTTAAGCGTAGTCTGGGACGTCGTATGGGTAGTCAATATTTTTTTGATCGACAATGAG |  |
| F1 | - pGCP123-*mprF1*-HA - pGCP123-*mprF2*-HA - pGCP123-*mprF2* (*D731A, R734S*)-HA - pGCP123-*mprF2* - pGCP123-*mprF2* (*D731A, R734S*) | gtaaaacgacggccagt | Screening for transformants containing pGCP123-mprF2  Same primers used for Sanger sequencing |
| R3 |  | acgctaaaacgtctcagaaac |  |
| pmprF2-MT-F | pGCP123-*mprF2* | ATGAAGTTGGTACCATCGCATTGATGAGCCACCATAAAGAAAAAGCCCC | Inverse PCR to create D731A, R734S mutation on *mprF2* |
| pmprF2-MT-R |  | TGGTACCAACTTCATTTGTGTAACTAGGAATGA |  |
| pmprF2-delHA_F | - pGCP123-*mprF2* (*D731A, R734S*)-HA - pGCP123-*mprF2* | ttaataaaaaatcgaTTATTAGTCAATATTTTTTTGATCGACAATGAGTAAAGCAATCATCACGTA | Inverse PCR to create remove HA affinity tag |
| pmprF2-delHA_F |  | tcgattttttattaaaacgtctcaaaatcgtttctg |  |
| pGCP123-P*srtA*_linear_F | pGCP123-PsrtA | tgcagcccgggggatccactagt | Inverse PCR to linearise the pGCP123-P*srtA* plasmid |
| pGCP123-P*srtA*_linear_R |  | aattcgatatcaagcttatcgataccgtcgacctcgagattctccc |  |
| 1. Insert-mprF1_F | genomic DNA | gcttgatatcgaattGAGATATGAAAGAGGCTATAAGATGAAAAAAAA | To create insert for ligation into pGCP123-P*srtA* linearized vector.  Primer pairs: (1+4), (1+2), (1+3) |
| 2. Insert_mprF1_(N-term)_R | genomic DNA | atcccccgggctgcaTTATTATCACCTAAAGTAACAGTTGTTTCGG |  |
| 3. Insert_mprF1_(C-term)_F | genomic DNA | gcttgatatcgaattATGGGGGTTTGAGGAAGCGCGGTT |  |
| 4. Insert-mprF1_R | genomic DNA | atcccccgggctgcaAGAGTTAAATTTCAGTCATTGCTTCT |  |
| 5. Insert_mprF2_F | genomic DNA | gcttgatatcgaattCTTAAAGGAAATGAAGGTGTCTAAATGAA | To create insert for ligation into pGCP123-P*srtA* linearized vector.  Primer pairs: (5+8), (5+6), (5+7) |
| 6. Insert_mprF2_(N-term)_R | genomic DNA | atcccccgggctgcaTTATTAAACTTGGTGTTTTTTCCCTTGGA |  |
| 7. Insert_mprF2_(C-term)_F | genomic DNA | gcttgatatcgaattATGGGTGAGTTTCCAGAAGATAGTATCTT |  |
| 8. Insert_mprF2_R | genomic DNA | atcccccgggctgcaTTTTGTTAGTCAATATTTTTTTGATCGACAAT |  |

**Table S1G.** Antimicrobial peptides used

| **Antimicrobial peptide** | **Concentrations (µg/ml)** | **Class** | **Net Charge (in physiological pH**) | **Manufacturer** |
| --- | --- | --- | --- | --- |
| Human β-defensin 2 (hBD-2) | 3.125, 6.25, 12.5, 25, 50 | Beta-defensin | +6 (Cationic) | Peptide Institute Inc., Japan |
| LL-37 | 3.125, 6.25, 12.5, 25, 50 | Cathelicidin | +6 (Cationic) | Peptide Institute Inc., Japan |
