## Supplementary Text 1A-B, Supplementary References for "Depleting cationic lipids involved in antimicrobial resistance drives adaptive lipid remodeling in *Enterococcus faecalis*"

**Supplementary Text 1A: TLC Spot Identification**

To identify spots in the 1D TLC in **Fig. 2A**, iodine and ninhydrin staining of 1D-TLCs of empty vector controls of wild type, ∆*mprF1*, ∆*mprF2*, ∆*mprF1*∆*mprF2* and p*mprF2*-HA complemented ∆*mprF2*, ∆*mprF1*∆*mprF2* together with PG, CL and lysyl-PG standards were done. **(Fig. S4A, B)**. 32P radiolabeled lipids of the same samples were also run as 1D TLCs **(Fig. S4C)**. In the iodine-stained TLC, 6 major spots were identified, labelled 1 to 6.

Spot 3’s migration position matches that of the PG and CL standards, suggesting that spot 3 might contain PG and CL that co-migrate together **(Fig. S4A)**. Spot 3 was also not strongly stained with ninhydrin and was present in the 32P radiolabeled samples **(Fig. S4B, C)**. This highly suggests that PG and CL comigrate together in spot 3 since neither PG and CL are amino modified, and have phosphate present in their glycerophosphate headgroup.

Spot 4’s migration position corresponds with L-PG and is also absent in empty vector controls of ∆*mprF2* and ∆*mprF1*∆*mprF2* **(Fig. S4A)**. Furthermore, spot 4 is stained with ninhydrin along with the L-PG control at the same location and this spot was also present in the 32P radiolabeled samples **(Fig. S4B, C)**. Taken together, this suggests that spot’s 4 identity is L-PG, since it is amino-modified, and phosphate is present in their glycerophosphate headgroup.

The other spots did not match the migration of the standards are spots 1, 2, 5 and 6. To assign identities to these spots, WT and ∆*mprF1*∆*mprF2* lipid extracts were run as 1D-TLCs under the same conditions and all spots were scraped off and lipids extracted for LC-MS/MS analysis **(Fig. S4D)**. The observed masses for each of the spots and identities are listed in **Excel Table S1C.** Spot 1 was found to contain DGDAG, spot 3 was confirmed to contain PG. CL however was unable to be detected in spot 3 due to the difficulty in ionizing CL for mass spectrometry. Spot 4 was confirmed to contain L-PG, spot 6 was confirmed to contain GPDGDAG, while Spot 5’s identity is likely to be a D-alanine modified GPDGDAG (D-ala-GPDGDAG) **(Excel Table S1C)**.

However, the identity of spot 2 is unclear from MS analysis. 18 unique species were identified from this spot that displayed 3 common product ions from MS2 analysis (m/z: 379.1, 397.1 and 415.1) suggesting that they belong to the same class. Fatty acyl product ions were also detected in MS2 suggestion that contents of this spot are indeed lipids. MS1 observed masses also do not match the theoretical exact masses of the other lipid classes that we expected – PG, LPG, Ala-PG, Arg-PG, DGDAG, MGDAG, GPDGDAG **(Excel Table S1C)**. This spot also does not stain with ninhydrin suggesting that this unknown class is not amino modified **(Fig. S4B).** Since spot 2 shows up in 32P labeled samples **(Fig. S4C, E)**, it is likely that it is a phosphorous containing lipid. Hence, with this information the spot 2’s identity has been assigned according to **Fig. S4D** where it is renamed “P-containing”.

To separate the co-migrating PG and CL spots, 2D-TLC was performed. Using correspond locations on 1D-TLC plates, spots on 2D-TLC plates can be assigned. As for the co-migrating PG and CL in spot 3, the distal spot was assigned as CL and the proximal spot PG since the solvent system in the second dimension is known to cause CL to migrate ahead of PG with theoretical Rf values of 0.38 for CL and 0.31 for PG (TLC Solvent Systems – Lipid Migration, Avanti Polar Lipids Inc.) **(Figure S4E)**. To verify that these two spots are truly PG and CL, 32P-labelling was carried out with 2D-TLC in the same solvent systems. Indeed, these 2 spots appear in the 32P-labelled samples as well.

In addition, three spots were also observed to be present in place of the L-PG spot in ∆*mprF2* and ∆*mprF1*∆*mprF2* in the 1D-TLC with 14C radiolabeled samples in **(Fig. 2A)**. These species are likely also present in the wild type, as they appear to overlap with the L-PG spot on 2D-TLCs of the wild type and ∆*mprF1*∆*mprF2* **(Fig 2E).**

**Supplementary Text 1B: Supplementary Methods**

**Bacterial Strains and Growth Conditions**

The wild type (WT) parental strain used was *E. faecalis* OG1RF (1) and *∆mprF1* and *∆mprF2* mutants used were described previously (2, 3). Unless otherwise stated, all bacterial strains were grown overnight to late stationary phase for 16-18 hours in their respective media at 37 ^o^C in static conditions and stored in 25% glycerol solution at -80 ^o^C when long-term storage was required. If mid-log phase cultures were required, late stationary phase cultures were sub-cultured 1:10 dilution into fresh media and grown at 37 ^o^C, 250 rpm shaking conditions until an OD_600_ of 0.5 ± 0.05 was reached. When normalisation of cultures was required, they were centrifuged at 6,000 rcf for 5 min at 4 ^o^C and cell pellets were washed twice with 1mL of phosphate buffered saline (PBS). Cell suspensions were then normalised to an OD_600_ of 0.7 or required optical density by diluting with PBS. All strains, their respective genotypes, sources, and growth media are listed in **Table S1A**.

**Molecular cloning**

To create pGCP123-*mprF1*-HA and pGCP123-*mprF2*-HA plasmids for expression of MprF with the hemagglutinin (HA) affinity tag, inserts comprising of *mprF1* and *mprF2* were generated via PCR

were used where the HA-tag sequence was incorporated into the reverse primer sequence. Linearized pGCP123 plasmids were generated by restriction digestion with PstI and EcoRI for ligation with *mprF1*-HA, and PstI and SphI for ligation with *mprF2*-HA. The linearised plasmids and inserts were ligated using the In-Fusion HD Cloning system (Takara, Japan) and transformed into Stellar™ Competent Cells (Takara, Japan) according to the manufacturer’s instructions. Plasmids were then extracted from the stellar competent cells using the Monarch^®^ Plasmid Miniprep Kit. They were then transformed into electrocompetent *E. faecalis* ∆*mprF1*, ∆*mprF2* and ∆*mprF1*∆*mprF2*. Expression was verified by SDS-PAGE and western immunoblot as described above. It was discovered that only the pGCP123-*mprF2*-HA plasmids displayed expression (data not shown).

To create the pGCP123-*mprF2* and pGCP123-*mprF2*(D731A, R734S) plasmids, site directed mutagenesis was performed on the pGCP123-mprF2-HA plasmid by inverse PCR to introduce the D731A, R734S mutation using Q5^®^ High-Fidelity DNA Polymerase (New England BioLabs, USA). The linearised PCR product was circularised using the In-Fusion HD Cloning system (Takara, Japan) and transformed into Stellar™ Competent Cells (Takara, Japan) according to the manufacturer’s instructions. Plasmids were then extracted from the stellar competent cells using the Monarch^®^ Plasmid Miniprep Kit. Inverse PCR was again performed to remove the hemagglutinin (HA) affinity tag and the rest of the steps repeated to introduce the plasmid into *E. coli*. Plasmids were then transformed into electrocompetent *E. faecalis* ∆*mprF2* and ∆*mprF1*∆*mprF2*.

To create p*mprF1*, p*mprF1-N-terminal*, p*mprF1-C-terminal*, p*mprF2*, p*mprF2-N-terminal*, p*mprF2-C-terminal* plasmids, inserts corresponding to either *mprF* and their individual domains were created using Q5^®^ High-Fidelity DNA Polymerase. Plasmids were extracted from *E. coli* DH5α harboring pGCP123-P*srtA* using the Monarch^®^ Plasmid Miniprep Kit. Inverse PCR was performed to obtain the linearized plasmid which was ligated using the In-Fusion HD Cloning system with the corresponding inserts and transformed into Stellar™ Competent Cells. Plasmids were then transformed into electrocompetent *E. faecalis* ∆*mprF1*∆*mprF2*.

Transformants were selected using by BHI or LB agar containing the appropriate antibiotics as listed in **Table S1A**. Screening of transformants was performed using colony PCR using Taq DNA Polymerase, recombinant (5 U/µL) (Thermo Scientific, USA) according to the manufacturer’s instructions. Gel electrophoresis to assess for product sizes was performed using 1% w/v agarose gel in TAE buffer ran at 100 V for 30 minutes followed by ethidium bromide staining for 10-15 minutes. After each transformation step, if plasmids passed the colony PCR check, they were extracted and sent for Sanger sequencing for their inserts to ensure the correct sequence was present (1st BASE DNA Sequencing Services, Singapore). PCR purification was performed using Wizard^®^ SV Gel and PCR Clean-Up System (Promega, USA) according to the manufacturer’s instructions. Primers used for the described PCR steps are shown in **Table S1F**.

**RNA isolation and RT-qPCR**

Real time quantitative-PCR (RT-qPCR) was performed on RNA isolated from WT and ∆*mprF1*∆*mprF2* strains for *pgsA* and fatty acid biosynthesis genes. Overnight cultures were subcultured 1:10 in BHI and grown to mid-log phase (OD_600_ 0.5 ± 0.05). RNA extraction and sample preparation for RT-qPCR were done as previously described with the following modifications (4). 1 mL of RNAprotect Bacteria Reagent (Qiagen, Germany) was added to 500 μL of mid-log phase culture and incubated for 5 minutes at room temperature. Cells were then pelleted and rinsed with PBS to thoroughly remove the RNAprotect reagent. Cell pellets were then resuspended in 20 mg/mL lysozyme (Sigma-Aldrich, USA) in lysis buffer (10 mM Tris-HCl pH 7.0, 1 mM EDTA, 50 mM NaCl and 0.74 M sucrose) and incubated for 1 hour. Cells were then pelleted and washed once with PBS. 1 mL of ice-cold TRIzol™ Reagent (Ambion, USA) was added and the cell pellet resuspended by pipetting action. 200 μL of ice-cold chloroform (Fisher, USA) was added to the tube and the suspension mixed by shaking gently before incubating on ice for 2 minutes. The mixture was then centrifuged for 15 minutes at 12,000 rcf at 4 ^o^C to induce phase separation. Next, 500 μL of the upper aqueous phase was carefully removed and added to the spin-column from the RNeasy Mini Kit (Qiagen, USA). Purification was then carried as specified by the manufacturer’s instructions and eluted in 40 μL of Nuclease-free water (Ambion, USA). RT-qPCR was next performed using the respective primers listed in **Table S1C** below on a StepOnePlus™ Real-Time PCR System (Applied Biosystems, USA). The product from cDNA synthesis run without template added was used as a negative control. Absolute quantification was carried out by running standard curves of the respective targets alongside on the same qPCR plate where the absolute quantities of each target in the sample were calculated using the StepOne™ software (Applied Biosystems, USA). If relative quantification was required, the delta-delta Ct method was used to calculate fold change values.

**RNA isolation and RNA sequencing**

Sequencing of RNA was done from OG1RF, *∆mprF1, ∆mprF2*, and *∆mprf1∆mprF2* strains. Bacteria cultures were grown overnight, statically in BHI media at 37^o^C. Starter cultures were subcultured 1:10 and grown to an optical density of 0.5 at 600 nm. Total RNA was extracted by the UltraClean^®^ Microbial RNA Isolation Kit (MO BIO Laboratories Inc (now Qiagen, Germany)), USA) from *E. faecalis* strains according to the manufacturer’s instructions. Extracted RNA samples were subjected to rigorous DNase treatment using TURBO DNA-free™ kit (Ambion^®^, USA) and subjected to ribosomal depletion with Ribo-Zero™ Magnetic Kits (Lucigen, USA) according to their respective manufacturer’s protocol. Quantification of RNA and DNA were performed using Qubit™ RNA Assay Kits and Qubit™ dsDNA HS Assay Kits (Invitrogen, USA), respectively. Integrity of RNA was analyzed by gel electrophoresis using Agilent RNA ScreenTape (Agilent Technologies, USA). cDNA synthesis was done using NEBNext^®^ RNA First Strand Synthesis Module and NEBNext^®^ Ultra Directional RNA Second Strand Synthesis Module (New England Biolabs, USA), and purified using AMPure XP beads (Beckman Coulter, USA). RNA library preparation was done by the sequencing facility of Singapore Centre of Life Science Engineering (SCELSE, Singapore) and RNA sequencing was done by the sequencing facility at Genome Institute of Singapore (GIS, Singapore) using HiSeq 2500.

**Growth kinetics (OD_600_ and CFU/mL)**

Overnight cultures were subcultured in Brain Heart Infusion (BHI) liquid broth at a 1:100 dilution. Optical density readings at a wavelength of 600 nm (OD_600_) were taken immediately (t= 0 hr) using a UV spectrophotometer (Shimadzu, Japan). Concurrently, 200 μL of culture was added to a 96-well microtitre plate (Thermo Fisher Scientific, USA) and serially diluted up to 10^-8^ using 1X sterile phosphate buffered saline (PBS). 5 μL of bacterial suspension from each dilution was then spotted 5 times onto a BHI agar plate, for a total of 5 technical replicates. This process was repeated every hour for the next 8 hours, and a final reading was taken after 24 hours. The spotted agar plates were then incubated statically overnight at 37 ^o^C and colony forming unit (CFU) enumeration was subsequently performed when the colonies were big enough to be counted.

**Growth kinetics (Plate reader based)**

Overnight cultures were normalised to OD_600_ of 0.7 and diluted 200-fold (for BHI) or 100-fold (for chemically defined media (CDM)) before inoculation in 200 μL of media in 96-well plates in a ratio of 1:25. The 96-well plates were incubated at 37 ^o^C in a Tecan Infinite^©^ M200 Pro spectrophotometer (Tecan, Switzerland) and absorbance was read at 600 nm every 10 minutes for 18 hours (for BHI) or 72 hours (for CDM). The components of the CDM used are shown in **Table S1D.** If fatty acid supplementation was required, fatty acids were prepared as 10 mg/mL stocks in ethanol for linoleic acid, oleic acid, *cis*-vaccenic acid, palmitoleic acid, stearic acid, palmitic acid, and lauric acid, and as 1 mg/mL in ethanol for arachidic acid.

**Live/Dead assay**

Overnight cultures were subcultured in BHI liquid broth at a 1:10 dilution and grown to mid-log phase (OD_600_ 0.5 ± 0.05). These bacterial cultures were washed once with PBS and normalised to OD_600_ 0.5. Viability staining was performed by incubating the cells with 1 μL of a SYTO9 and propidium iodide (PI) live dead stain mix (LIVE/DEAD® *Bac*Light™Bacterial Viability Kit, Life Technologies, USA) for 15 minutes in the dark at room temperature. Stained cells were then washed with 200 μL of 0.01M low-salt phosphate buffer (PB). Wet mounting was done by smearing 10 μL of the sample onto clear 1.0-1.2mm microscope slides (Biomedia, Singapore) and covered with a square 22 x 22 mm coverslip. Slides were then imaged with an inverted epi–fluorescence microscope (Zeiss Axio observer Z1, Carl Zeiss GmbH, Germany) fitted with an 100X Oil immersion objective (numerical aperture 1.4, optovar 1.6X), an AF488/FITC filter cube (460-490 nm band pass excitation filter, 515-550nm band pass barrier filter) and an AF568/Cy3 filter cube (530-550nm band pass excitation filter, 590nm long pass barrier filter). Phase, fluorescent and merged field images were captured using AxioVision software and saved for computational analysis.

**MS analysis of TLC spots**

TLC spots to be identified were scraped and silica fragments were collected in Eppendorf tubes. Lipid extraction was carried out as described in the main methods section. Chromatography separation was achieved by hydrophilic interaction liquid chromatography (HILIC) on a Phenomenex Kinetex HILIC column (150 x 2.10 mm, 2.6μM, 100Å), using Vanquish LC system (ThermoScientific). The column temperature was 40 °C, the autosampler was kept at 10 °C, and 1 μL of sample was injected. Solvent A was acetonitrile/25mM aqueous ammonium formate pH4.6 (1/1 v/v), solvent B was acetonitrile/25mM aqueous ammonium formate pH4.6 (95/5 v/v. Gradient elution started at 1 % solvent A, increased linearly to 75% A in 6 min, then increased linearly to 90 % A in 1 min, then decreased back at 1 % A in 0.1 min, and held for 3 min (total runtime 10.1 min). The flow rate was 500μL/min. The column effluent was introduced into a Thermo QExactive plus mass spectrometer.

FT-MS spectra in the range of m/z 250-1250 were acquired in profile mode at a resolution setting of 140,000 (FWHM at m/z 200) with an AGC setting of 3x106 and maximum ion time of 200 ms. Product ion scans in centroid mode were acquired in data-dependent analysis mode, with a resolution setting of 17500 (FWHM at m/z 200), AGC setting of 1x105, normalized collision energy of 25, and isolation window of 1.4 m/z, and fixed first mass m/z 80. Samples were analyzed in positive and negative ionization.
